# Coupling of tubulin acetylation to microtubule stabilization through molecular mimicry

**DOI:** 10.64898/2026.07.08.736797

**Authors:** Juan M. Perez-Bertoldi, Thanh mai Julie Dang, Mert Golcuk, Eva Lanska, Danilo Lopes, Veronique Henriot, Jingyi Luo, Rui Zhang, Shih-Chieh Ti, Carsten Janke, Zdenek Lansky, Mert Gur, Filippo Del Bene, Eva Nogales

**Affiliations:** Biophysics Graduate Group, University of California, Berkeley, CA, USA; Sorbonne Université, INSERM, CNRS, Institut de la Vision, Paris 75012, France; Department of Computational and Systems Biology, University of Pittsburgh, Pittsburgh, PA, USA; Institute of Biotechnology, Czech Academy of Sciences, BIOCEV, Vestec, Czech Republic; Institut Curie, Université PSL, CNRS UMR3348, INSERM U1368, Orsay, France; Université Paris-Saclay, CNRS UMR3348, INSERM U1368, Orsay, France; School of Biomedical Sciences, Faculty of Medicine, The University of Hong Kong, Hong Kong SAR, China; Department of Biochemistry and Molecular Biophysics, Washington University in St. Louis, School of Medicine, St. Louis, MO, USA; Department of Molecular and Cell Biology, University of California, Berkeley, CA, USA; Molecular Biophysics and Integrative Bioimaging Division, Lawrence Berkeley National Laboratory, Berkeley, CA, USA; Howard Hughes Medical Institute, University of California, Berkeley, CA, USA

## Abstract

Microtubules support diverse cellular functions through regulation by microtubule-associated proteins and tubulin post-translational modification, yet how these two layers are mechanistically integrated remains unclear. α-tubulin acetylation marks mechanically resilient microtubules, and its incorporation in defined microtubule sub-populations is not well understood. Here, we identify MTCL1 as a molecular link between microtubule stabilization and post-translational modification installation. We find that MTCL1 stabilizes microtubules and alters the luminal surface when copolymerized with tubulin, remodeling α-tubulin and enhancing αTAT-mediated tubulin acetylation through molecular mimicry. This effect depends on assembly history and is not observed in pre-assembled microtubules. Targeted deletion of MTCL1 in zebrafish impacts axonal organization, leading to motor defects and increased seizure susceptibility. These findings establish MTCL1 as a licensing factor that couples microtubule stabilization with acetylation to regulate neuronal function.

## Introduction

Microtubules (MTs) are essential cytoskeletal polymers, built from repeating α/β-tubulin heterodimers, that organize the cell interior, drive cell division, support cargo transport, and help shape cell morphology. To fulfill their many cellular functions, MT biophysical properties are locally regulated by the action of molecular motors, the binding of non-motile MT-associated proteins (MAPs)^1^, and by a rich “tubulin code” of post-translational modifications (PTMs)^2^. Together, these control layers allow cells to assemble MTs with distinct dynamics, binding specificities, and array geometries that support a large number of physiological functions. Within this complex regulatory landscape, MT cross-linking factor 1 (MTCL1) is essential for cellular processes in which MTs are nucleated away from the centrosome. These include the formation of cortical bundles in epithelia, where this MAP was discovered^3^, the perinuclear network that supports Golgi function^4^, and ordered neuronal fascicles where MTCL1 contributes to establishing the axon initial segment (AIS), a boundary and selective barrier between soma and axon^5^. MTCL1 contains a large, predicted coiled-coil scaffold and two distinct MT-binding regions located at its termini. Cellular and *in vivo* studies have shown that the N-terminal MT-binding domain (MTCL1^N-MTBD^) drives cross-linking and MT bundle formation, whereas the C-terminal domain (MTCL1^C-MTBD^) stabilizes MTs and its presence correlates with increased acetylation levels^3,6^.

Disruption of MTCL1 function has been linked to neuronal dysfunction and human disease. Mutations associated with spinocerebellar ataxia map to the MTCL1 C-terminal region, and additional truncating variants lacking this domain have been identified^5,7^. In mice, loss of MTCL1 in cerebellar Purkinje cells leads to disorganization of AIS-associated MT bundles, mislocalisation of ankyrin G, and locomotor defects^5^. These observations suggest that proper organization and stabilization of MT arrays by MTCL1 is critical for neuronal integrity.

Tubulin PTMs fine-tune the mechanical properties and MAP binding of specific MT populations to regulate their cellular functions^2^. While most of the well-characterized PTMs are present on the flexible, surface exposed C-terminal tails of α- and β-tubulin, acetylation of α-tubulin on Lys40 (K40Ac) is localized on a flexible loop facing the MT lumen^8–10^. In human cells, acetylation of tubulin K40 is carried out by the acetyltransferase αTAT1/MEC-17^11,12^. αTAT1 needs to gain access to the MT lumen through open ends or transient lattice defects^13–16^ and then engage the H1–S2 internal loop of α-tubulin containing K40.

K40Ac accumulates in long-lived MTs and modulates mechanical behavior, providing increased resilience to repeated bending and compression^17–19^. Tubulin acetylation is highly conserved and its misregulation is implicated in disease, from cancer^20,21^ to ciliopathy-related defects in tissue morphogenesis^12,22,23^ and neurodevelopmental/neurodegenerative phenotypes^24–28^. Particularly, alterations in tubulin acetylation are broadly implicated in neuropathies, from axonal degeneration in Charcot-Marie-Tooth disease^29^, to neurodegenerative diseases such as Alzheimer’s and Parkinson’s. Tubulin hyperacetylation has been associated with early Lewy body pathology and α-synuclein aggregation in Parkinson’s^30,31^, and documented in familial forms of both diseases^32–34^.

Interestingly, multiple studies report that MT arrays enriched in MTCL1 also show higher levels of K40 acetylation, and that loss of MTCL1 reduces local acetyl-K40^4–6^. What remains unclear is whether MTCL1 binding to MTs directly drives increased acetylation, or instead reflects the well-established correlation between MT stability and acetylation^10^, or a more complex coupling between MT bundling, lattice mechanics, and enzyme accessibility.

In this study, we combined biochemical reconstitution, cryoelectron microscopy (cryo-EM), all-atom molecular dynamics (MD) simulations, fluorescence microscopy, and both cell-based, and *in vitro* functional assays to define how MTCL1^C-MTBD^ interacts with MTs and connect these molecular insights to their physiological function by examining AIS organization in genome-edited zebrafish.

## Results

### MTCL1^C-MTBD^ reinforces β-tubulin lateral contacts to stabilize MTs

To better understand MTCL1-MT interactions, we recombinantly expressed and purified human MTCL1^C-MTBD^ (Figure 1a, Supplementary Figure 1a) and used cryo-EM to visualize how this domain binds MTs. In our initial approach, MTs were pre-assembled and stabilized with paclitaxel (Taxol), after which MTCL1^C-MTBD^ was added prior to vitrification (Figure 1b). Using a MT-tailored image processing workflow that included explicit seam detection for each filament, helical reconstruction, and symmetry expansion^35^, we obtained a 2.7 Å reconstruction of MTCL1^C-MTBD^ on Taxol MTs (Figure 1 c-d, Supplementary Figures 2 and 3a). The map shows a segment of 20 residues, from 1739 to 1758, of MTCL1^C-MTBD^ binding MTs at the inter-protofilament groove, where it forms tight contacts with neighboring β-tubulins and reinforces lateral tubulin interactions (Figure 1d).

**Figure 1:**
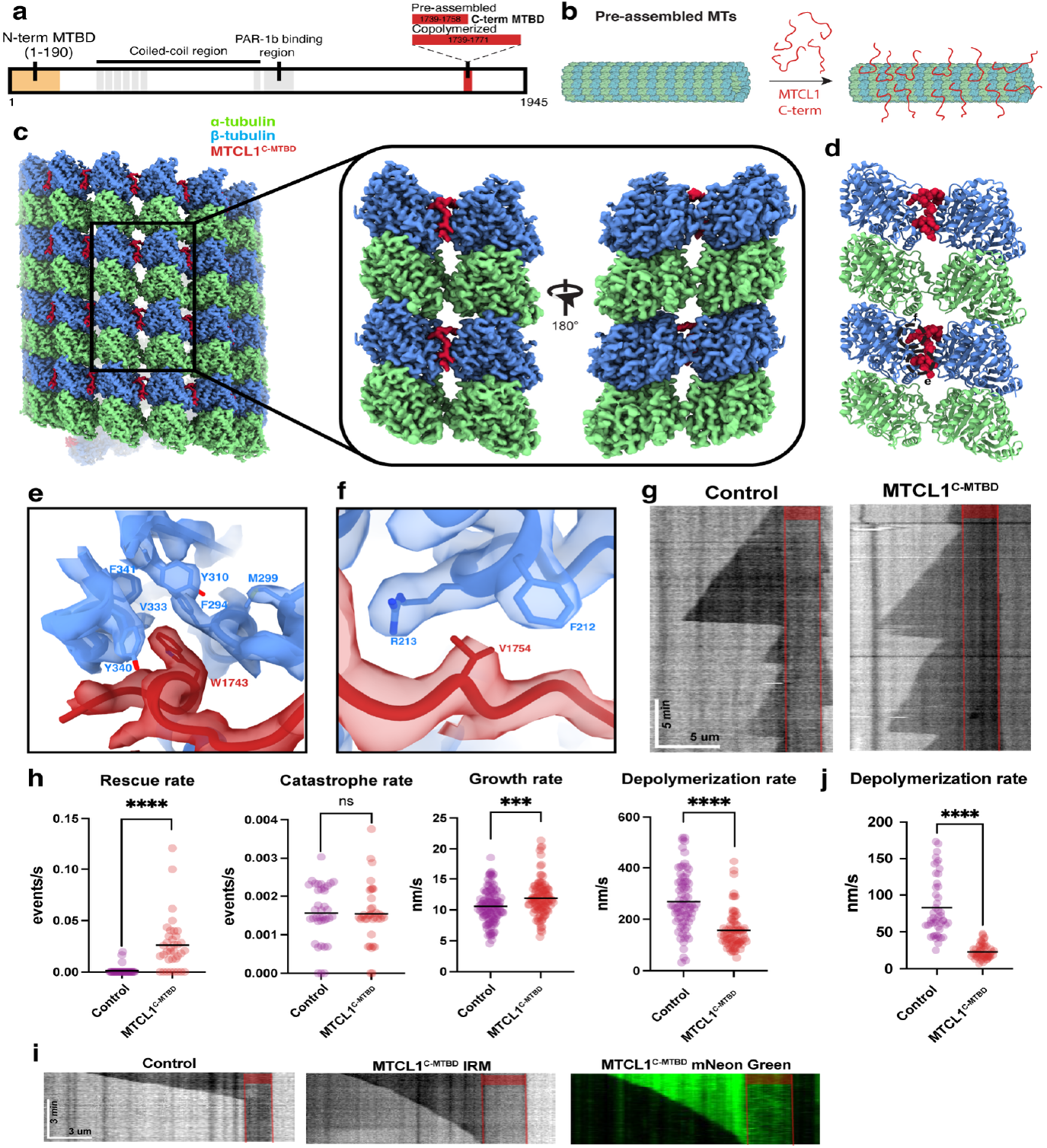
MTCL1^C-MTBD^ is a MT stabilizer. **(a)** Domain organization of human MTCL1. **(b)** Schematic showing strategy for sample preparation of pre-assembled Taxol-stabilized MTs + MTCL1^C-MTBD^ used for cryo-EM structure determination. **(c)** Cryo-EM reconstruction of MTCL1^C-MTBD^-decorated MT under post-assembly binding conditions. The inset shows four tubulin dimers across an interprotofilament interface, in outside (left) and luminal (right) views, with MTCL1^C-MTBD^ binding at the lateral interface. α-tubulin, β-tubulin and MTCL1^C-MTBD^ are shown in light green, cornflower blue and fire brick, respectively. **(d)** Ribbon representation of the two adjacent protofilaments shown on the left side of the inset in c, showing MTCL1^C-MTBD^ in sphere representation binding at the inter-protofilament groove. Regions of interest are encircled with dashed lines and expanded in panels **e** and **f. (e)** Close-up view of the map and fitted model showing MTCL1^C-MTBD^ residue W1743 engaging an aromatic pocket on β-tubulin. **(f)** Close-up view of the map and fitted model showing MTCL1^C-MTBD^ residue V1754 near β-tubulin residues F212 and R213. **(g)** Representative kymographs of dynamic MTs under control conditions (left) and in the presence of MTCL1^C-MTBD^ (right), visualized by interference reflection microscopy (IRM). Horizontal and vertical scale bars represent 5 µm and 5 min, respectively. **(h)** Quantification of MT dynamic parameters, including catastrophe and rescue frequencies and growth and depolymerization rates, under control conditions and with MTCL1^C-MTBD^. Data was obtained from a total of 48 MTs (control) and 42 MTs (MTCL1), pooled from three independent experiments. **(i)** Representative kymographs of MTs after washout under control conditions (left) or in the presence of MTCL1 (middle), visualized by IRM. The corresponding fluorescence kymograph of MTCL1^C-MTBD^ bound to the same MT shown in the middle panel was acquired using 488-nm excitation (right). **(j)** Quantification of MT depolymerization rates after washout under control conditions and with MTCL1^C-MTBD^. Data was obtained from a total of 40 MTs (control) and 39 MTs (MTCL1^C-MTBD^), pooled from three independent experiments. In all quantification plots, data are shown for individual MTs, with the mean indicated. Red boxes denote GMPCPP-stabilized MT seeds. Statistical significance was assessed using unpaired two-tailed t-tests. ***P < 0.001; ****P < 0.0001.

Among the MTCL1^C-MTBD^-tubulin interactions, a conserved tryptophan W1743 provides a prominent anchor on one of the protofilaments by inserting into an aromatic pocket on β-tubulin (Figure 1e, Supplementary Figure 4). In α-tubulin, the equivalent pocket contains instead polar or bulky side chains (Supplementary Figure 5), explaining why MTCL1^C-MTBD^ binding is not observed at α-α interfaces. Additionally, V1754 packs against β-tubulin F212 and the methylene portion of R213 at the lateral contact, forming favorable van der Waals interactions (Figure 1f). V1754 also contributes to β-tubulin selectivity, as in α-tubulin, the position corresponding to β-tubulin’s F212 is an arginine (Supplementary Figure 5a). A patient-derived variant in a spinocerebellar ataxia allele, in which V1754 is replaced by methionine, was previously identified and assayed for MT stabilization in cells^5^. In our structure, the larger methionine side chain would not fit easily within the F212/R213 pocket, providing a structural basis for the reduced MT decoration reported for this mutant (Figure 1f).

The V1754M variant has been linked to lower acetylation readouts in cells^5^. While the most C-terminal density of MTCL1^C-MTBD^ extends towards the inter-protofilament fenestrations (Figure 1c-f), the structure did not show density in the MT lumen.

Given the expected stabilization of MTs by the fragment we characterized structurally, we reconstituted dynamic MTs *in vitro* and imaged their dynamics using total internal reflection fluorescence (TIRF) microscopy in the presence of purified MTCL1^C-MTBD^. Kymographs revealed that MTCL1 markedly stabilized MTs, primarily by increasing the rescue frequency, reducing the depolymerization rate, and by modestly increasing the growth rate, with no significant effect on the catastrophe frequency (Figure 1g, h). Thus, MTCL1^C-MTBD^ leads to an overall stabilization of dynamic MTs by strongly suppressing MT shrinkage. These results are consistent with our structural observations, which indicate that MTCL1 reinforces lateral contacts between protofilaments. To further test whether MTCL1 directly protects the polymer from disassembly, we performed a wash-out assay in which MTs were first polymerized with GTP in the absence of chemical stabilizers, either with or without MTCL1^C-MTBD^, and subsequently perfused with buffer lacking both tubulin and MTCL1. Under these conditions, undecorated MTs rapidly depolymerized, whereas MTCL1-decorated MTs exhibited a significantly reduced depolymerization rate (Figure 1i, j). Importantly, MTCL1 remained bound to the lattice after wash-out, supporting the notion that MTCL1 wedges between protofilaments to reinforce lattice integrity. *In vitro* co-sedimentation assays further indicated that MTCL1^C-MTBD^ can lower the critical concentration for spontaneous MT nucleation, which is relevant for this protein’s role in organizing centrosome-independent MT networks (Supplementary Figure 1b).

### MTCL1^C-MTBD^ extends into the MT lumen when copolymerized with tubulin

Many MAPs and motors bind to pre-formed MTs to stabilize the lattice, recruit other factors, remodel lattice architecture, generate force, or transport cargo. Additionally, MAPs can co-assemble with tubulin as the polymer is growing^36–39^. Many of these copolymerization-competent MAPs, including Tau, MAP6, and DCX, are enriched in the brain, where they contribute to neuronal MT organization. Given the neuronal localization of MTCL1, we examined human MTCL1 under conditions of tubulin copolymerization *in vitro*.

Taking advantage of its intrinsic stabilizing activity, we co-polymerized tubulin with MTCL1^C-MTBD^ in the absence of stabilizing drugs and determined a 2.7 Å cryo-EM structure (Figure 2a). The reconstruction reveals a pronounced extension of the MTCL1^C-MTBD^ density that inserts through the inter-protofilament fenestrations and into the MT lumen (residues 1759-1771), where it contacts α-tubulin of the neighboring dimer across the longitudinal interface. This configuration staples the inter-dimer junction and compacts the lattice by ∼1 Å with respect to GDP-lattice of undecorated MTs^40^, while preserving the contacts that reinforce the lateral interface between adjacent β-tubulins (Figure 2a, b). Together, these features further rationalize the robust stabilizing activity of MTCL1^C-MTBD^ on MTs (Figure 1g-j). To determine whether this extended MTCL1^C-MTBD^ binding mode was induced by copolymerization or by the absence of stabilizing drugs, we collected a cryo-EM dataset from a sample where MTCL1^C-MTBD^ was co-assembled with tubulin in the presence of Taxol. A systematic comparison of the population distribution after 3D classification of the different datasets showed that MTCL1^C-MTBD^ can adopt an extended footprint on MTs regardless of Taxol addition, suggesting that copolymerization is the primary mechanism enabling this class to be populated (Supplementary Figure 6).

**Figure 2:**
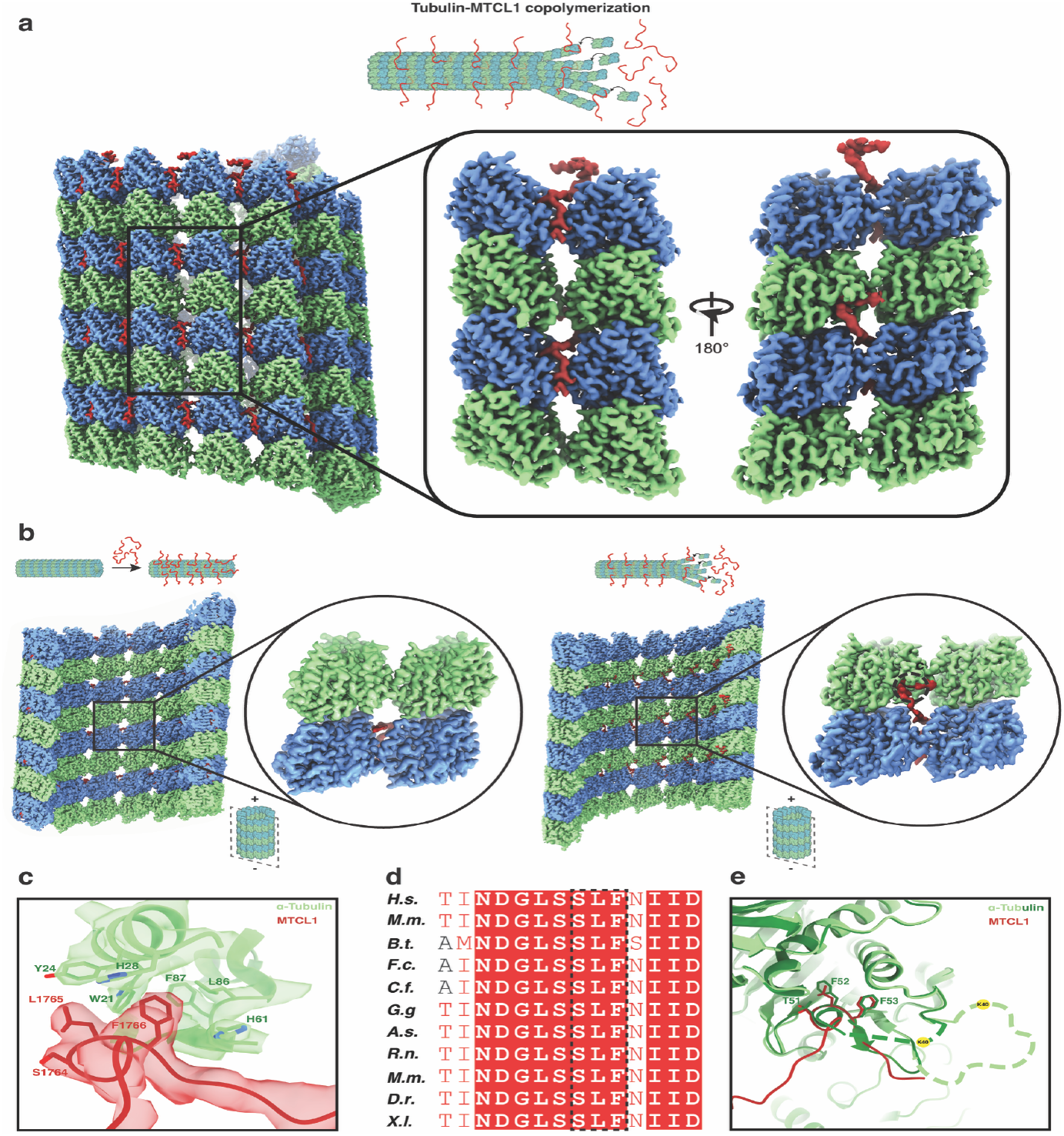
MTCL1^C-MTBD^ accesses the MT lumen under tubulin–MTCL1 copolymerization conditions. **(a)** (Top) Strategy for cryo-EM sample preparation under tubulin and MTCL1 copolymerization conditions. (Bottom) cryo-EM reconstruction of a MT copolymerized with MTCL1^C-MTBD^. The inset shows the outside (left) and luminal (right) views of the final, refined map, which include additional MTCL1^C-MTBD^ density extending into the MT lumen. Only two MTCL1 molecules are shown for simplicity. **(b)** Comparison of reconstructions from samples with MTCL1^C-MTBD^ added after or during MT polymerization, with insets highlighting absence or presence of MTCL1^C-MTBD^ luminal densities at an inter-dimer, inter-protofilament interface. **(c)** Cryo-EM map and fitted model for the conserved SLF motif in MTCL1^C-MTBD^ interacting with an α-tubulin hydrophobic pocket. **(d)** Multiple-sequence alignment around the SLF motif across representative species. **(e)** Ribbon diagram for α-tubulin and the SLF motif (red) as it displaces part of the H1-S2 loop in α-tubulin. α-tubulin in the presence and absence of MTCL1 is shown in light and dark green, respectively. The extension of the disordered part of the H1-S2 in each scenario is shown schematically with dashed lines. The position of K40 is shown with a yellow circle.

Modeling MTCL1^C-MTBD^ into the extended density identified a prominent luminal contact on α-tubulin involving a highly conserved Ser-Leu-Phe (SLF) motif comprising S1764, L1765 and F1766. S1764 forms a hydrogen bond with the backbone carbonyl of α-tubulin L242 (not shown), while L1765 and F1766 insert into a hydrophobic pocket on α-tubulin located adjacent to K40 within the H1–S2 loop (Figure 2c, d). Remarkably, the backbone geometry and side-chain chemistry of the SLF motif closely mimic those of α-tubulin residues T51, F52 and F53 that normally occupy this pocket but are not seen in our structure, revealing a striking case of molecular mimicry by MTCL1^C-MTBD^. In our model, the SLF motif displaces the α-tubulin residues and disrupts the ordered segment of the H1–S2 loop spanning residues 47–59, which becomes disordered and unresolved in the density. As a consequence, the K40-containing internal loop extends from a 10-residue disordered region (P37–D47) to a 22-residue disordered region (P37–G59) in the presence of MTCL1^C-MTBD^ (Figure 2e). We hypothesized that this increase in loop length and flexibility would enhance the accessibility of the H1–S2 loop to the tubulin acetyltransferase, thus providing a mechanism by which MTCL1^C-MTBD^ could directly modulate K40 acetylation on MTs.

### MTCL1^C-MTBD^ enhances α-tubulin K40 acetylation

To test our structure-based hypothesis that MTCL1^C-MTBD^ enhances tubulin acetylation by increasing the accessibility of the K40-containing H1–S2 loop, we established an *in vitro* acetylation assay using C. elegans αTAT2, a close structural homolog of human αTAT1 (Supplementary figure 7). We assembled MTs using recombinant, non-acetylated α1Aβ4B tubulin polymerized either in the presence or absence of MTCL1^C-MTBD^. To directly assess whether displacement of the H1–S2 loop is required for this putative enhancement, we also generated an MTCL1^C-MTBD^ variant in which the SLF motif was mutated to AKR (MTCL1^C-MTBD^ AKR) (Supplementary figure 1a). This substitution should disrupt MTCL1 luminal interactions that cause H1-S2 loop remodeling while presumably preserving outer-surface binding and MT stabilization. The unacetylated MTs were incubated with αTAT2 and acetyl-CoA, and acetylation was monitored over time by anti-Ac-K40 immunoblotting (Figure 3a).

**Figure 3:**
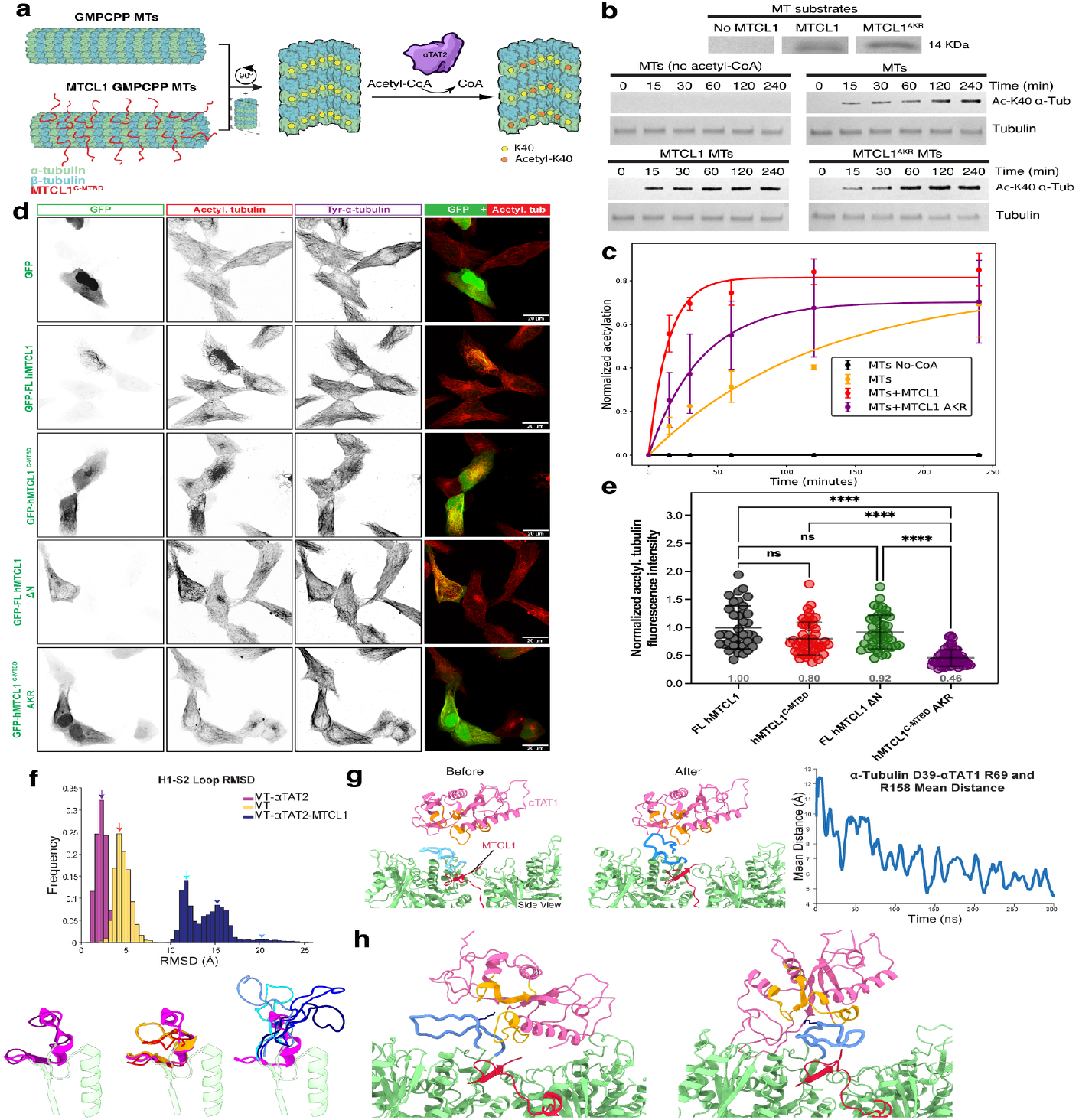
MTCL1^C-MTBD^ enhances α-tubulin K40 acetylation. **(a)** Schematic of the acetylation assay using non-acetylated α1Aβ4B MTs polymerized in the presence or absence of MTCL1^C-MTBD^ and incubated with αTAT2 and acetyl-CoA. **(b)** Coomassie Blue stained gel bands showing presence or absence of MTCL1^C-MTBD^/MTCL1^C-MTBD^ AKR in the MT substrates used in the assay (top). Time series of anti–Ac-K40 immunoblots for MTs (no acetyl-coA), MTs, MTs with MTCL1^C-MTBD^ and MTs with MTCL1^C-MTBD^ AKR (bottom). **(c)** Quantified kinetics of b. Curves for MTs (no acetyl-coA) (KAc = 0 min^-1^), MTs (KAc = 0.016 ± 0.005 min^-1^), MTs with MTCL1^C-MTBD^ (KAc = 0.14 ± 0.02 min^-1^) and MTs with MTCL1^C-MTBD^ AKR (KAc = 0.053 ± 0.003 min^-1^) are shown in blue, orange, green and red, respectively. The center line and whiskers represent the mean normalized acetylation and standard error (SE), respectively. **(d)** Representative immunofluorescence images of HeLa cells transfected for 24h with the indicated GFP-tagged human MTCL1 plasmid. Cells with lower expression levels are depicted, which were stained with antibodies against acetylated α-tubulin and tyrosinated tubulin. GFP transfection was used as control. Scale bar, 20 μm. **(e)** Quantification of the immunofluorescence in d. Fluorescence intensity ratios between acetylated/tyrosinated α-tubulin. Values were normalized to the average level of FL hMTCL1 cells. Each dot represents an individual cell; pool of three independent experiments (mean ± SD). Cells analyzed: GFP-FL hMTCL1, n = 38 cells; GFP-hMTCL1C-MTBD, n = 49 cells; GFP-FL hMTCL1 ΔN, n = 44 cells; GFP-hMTCL1C-MTBD AKR, n = 53 cells. Statistical significance was assessed using one-way ANOVA and the Kruskal-Wallis test. ns-not significant, ****P ≤ 0.0001. **(f)** (Top) Distribution of Cα-based RMSD values for the H1–S2 loop in three MD simulation conditions: αTAT2–bound MT, apo-MT after αTAT2 removal, and MTCL1^C-MTBD^–bound MT. RMSD values were computed relative to the ordered H1–S2 loop conformation in the αTAT2-bound MT structure (PDB:8Y9F). Arrows indicate the peaks of the RMSD distributions and the corresponding RMSD values from which the representative H1–S2 loop conformations shown in the bottom panel were selected. The colors of the RMSD distributions match those of the corresponding conformations shown below. (Bottom) The panel shows the H1–S2 loop conformations sampled from the RMSD peaks in the top panel, corresponding to αTAT2-bound MT (purple), apo-MT (orange), and MTCL1^C-MTBD^–bound MT (cyan, blue, and light blue), superimposed onto the αTAT2-bound MT structure (magenta). **(g)** Examples of H1–S2 loop conformations before and after αTAT1 was positioned at a fixed distance of 10 Å from tubulin, illustrating extension of the loop toward αTAT1 and formation of contacts with the enzyme (left and middle, respectively). Time evolution of the H1–S2 loop distance to αTAT1. The distance was calculated as the average of the Cα–Cα distances between α-tubulin Asp39 and αTAT1 Arg69, and between Asp39 and Arg158 (right). **(h)** Bound conformations of αTAT1 on the MT surface in the MTCL1^C-MTBD^– tubulin–αTAT1 ternary complex from two separate MD runs (final conformations), showing attachment of αTAT1 and placement of the H1–S2 loop/K40 into the αTAT1 active-site cavity. In g and h, residues corresponding to the αTAT1 active site are shown in orange.

Control reactions lacking acetyl-CoA showed no signal, confirming assay specificity (K_Ac_ = 0 min^-1^). MTCL1^C-MTBD^-bound MTs displayed a faster rise in Ac-K40 signal and reached a higher plateau compared to MTs alone within the same time window, pointing to a direct enhancement of acetylation by MTCL1^C-MTBD^ (Figure 3b). Quantification and single-exponential fitting revealed a larger apparent acetylation rate constant for MTCL1^C-MTBD^ bound MTs (KAc = 0.14 ± 0.02 min^-1^) compared with MTs alone (KAc = 0.016 ± 0.005 min^-1^). Consistent with our hypothesis, MTs polymerized in the presence of the mutant MTCL1^C-MTBD^ AKR exhibited an intermediate behavior (KAc = 0.053 ± 0.003 min^-1^). Thus, preventing SLF-dependent MTCL1 luminal engagement dampens the acceleration of K40 acetylation (Figure 3c).

Next, to assess the impact of MTCL1^C-MTBD^ on α-tubulin K40 acetylation in cells, we analyzed acetylation levels in HeLa cells expressing GFP-labeled wild-type (WT) hMTCL1^C-MTBD^, hMTCL1^C-MTBD^ AKR, hMTCL1 full-length (FL), or FL lacking the N-terminal MTBD (FL hMTCL1 ΔN). At high expression levels (as indicated by strong GFP fluorescence), all MTCL1 constructs similarly induced pronounced MT bundling that correlated with elevated acetylation levels (Supplementary Figure 8). To determine whether this strong expression and bundling behavior could be masking subtle but real differences in acetylation levels between the different MTCL1 constructs, we examined cells with lower expression of MTCL1. Under these conditions, the levels of K40 acetylation were markedly lower in cells expressing the mutant MTCL1^C-MTBD^ AKR when compared with the WT hMTCL1^C-MTBD^, FL hMTCL1, and FL hMTCL1 ΔN (Figure 3d, e), which is consistent with the *in vitro* findings.

Thus, MTCL1^C-MTBD^ enhances the efficiency and extent of α-tubulin K40 acetylation both *in vitro* and in cells. This functional effect is consistent with our structural observation that MTCL1^C-MTBD^ remodels the H1–S2 loop and our model that it increases its accessibility, providing a direct mechanistic link between MT binding by MTCL1^C-MTBD^ and accelerated K40 acetylation.

### MTCL1^C-MTBD^ modifies H1–S2 loop flexibility and accessibility to facilitate αTAT binding and K40 acetylation

To gain insight into how MTCL1^C-MTBD^ affects the H1–S2 loop dynamics, accessibility of human αTAT1, and subsequent acetylation of K40, we performed five sets of extensive allatom MD simulations. First, MD trajectories were initiated from the *C. Elegans* αTAT2–tubulin complex structure (PDB ID: 8Y9F), in which αTAT2 is bound to the MT and the K40-containing loop is engaged with the active site of the enzyme. The H1–S2 loop displayed low conformational fluctuations, consistent with its stabilization by αTAT2 binding (Supplementary Video 1). Next, MD simulations were initiated from the αTAT2–tubulin structure after removing αTAT2. The H1–S2 loop rapidly became largely unstructured, as reflected by increased Cα-based root-meansquare deviation (RMSD) and residue fluctuation values (Figure 3f, Supplementary Figure 9a, b and Supplementary Video 2), highlighting a close coupling between the structural state of the loop and αTAT2 binding. We next investigated the effect of MTCL1^C-MTBD^ binding on the H1-S2 loop, by first constructing a loop model of the regions not visible in the cryo-EM density due to disorder using our published protocol^41,42^ and subsequently performing MD simulations (Supplementary Video 3). In the presence of MTCL1^C-MTBD^, there was a pronounced increase in the loop’s conformational flexibility, resulting in higher solvent exposure and accessibility compared with the already elevated levels observed in tubulin alone (i.e. upon αTAT2 removal) (Figure 3f, Supplementary Figure 9a, b).

We next tested whether this more flexible state of the H1–S2 loop could influence recruitment of human αTAT1. We kept αTAT1 10 Å away from tubulin and observed the H1–S2 loop extending toward αTAT1 and effectively “catching” the enzyme (Figure 3g, Supplementary Video 4). Next, we allowed the enzyme to move freely and run seven independent MD simulation runs (2.1 µs total), initiated from distinct H1–S2 loop starting conformations (Supplementary Figure 9c, Supplementary Video 5). Association of αTAT1 with the MT luminal surface was observed in most trajectories (Figure 3h and Supplementary Figure 9d), and in two of these simulations H1–S2/K40 entered the αTAT1 active-site cavity (Figure 3h). Although the overall bound geometry differed among trajectories, αTAT-1 association was consistently accompanied by direct contacts with the H1–S2 loop, indicating that this loop provides a “capture” interaction that likely facilitates recruitment and the productive positioning of K40. These MD results further underscore the importance of the MTCL1^C-MTBD^-boosted H1–S2 loop flexibility and show that it allows it to explore an ensemble of states that can capture αTAT1, and position K40 at the active site for acetylation.

Overall, our MD simulations support and extend our structural, biochemical, and cellular findings, revealing that MTCL1^C-MTBD^-driven remodeling increases H1-S2 loop flexibility and accessibility for αTAT binding, thereby providing a mechanistic explanation for the enhanced K40 acetylation observed both *in vitro* and in cells.

### The C-terminal MTBD is required for MTCL1 localisation at the AIS *in vivo*

Having established the structural basis of MTCL1^C-MTBD^-MT engagement, we next asked how it contributes to MTCL1 function *in vivo*. To this end, we examined its role in targeting the protein to the AIS in neurons. We focused on caudal primary (CaP) motor neurons in the zebrafish spinal cord, a well-defined and extensively characterized neuronal population^43^. To ensure developmental consistency, analyses were restricted to a defined rostro–caudal region of the spinal cord, taking advantage of the stereotyped maturation gradient in zebrafish to compare neurons at equivalent stages.

The expression of fluorescently-tagged human MTCL1 constructs, either full-length (hMTCL1-FL) or lacking the C-terminal MTBD (hMTCL1-ΔC), enabled the live visualization and comparison of the protein subcellular localization in these neurons *in vivo* at 2 days post fertilization (dpf) (see methods, Figure 4a). hMTCL1-FL displayed a clear enrichment at the axon hillock and presumptive AIS, occupying on average (10.96 ± 1.25)% of the axon length. In contrast, deletion of the C-terminal MTBD abolished this spatial restriction, resulting in a broader distribution along the axon, spanning (39.5 ± 7.30)% of its length (p = 0.0005) (Figure 4b-d). A similar effect was observed in the soma: hMTCL1-FL was confined to a narrow region proximal to the AIS, covering (9.65 ± 2.49)% of the somatic area, whereas hMTCL1-ΔC extended over a significantly larger fraction (31.75 ± 5.12)% (p = 0.0006) (Figure 4b, c, e). These results show that the C-MTBD is essential for the spatial restriction of MTCL1 to the AIS *in vivo*.

**Figure 4:**
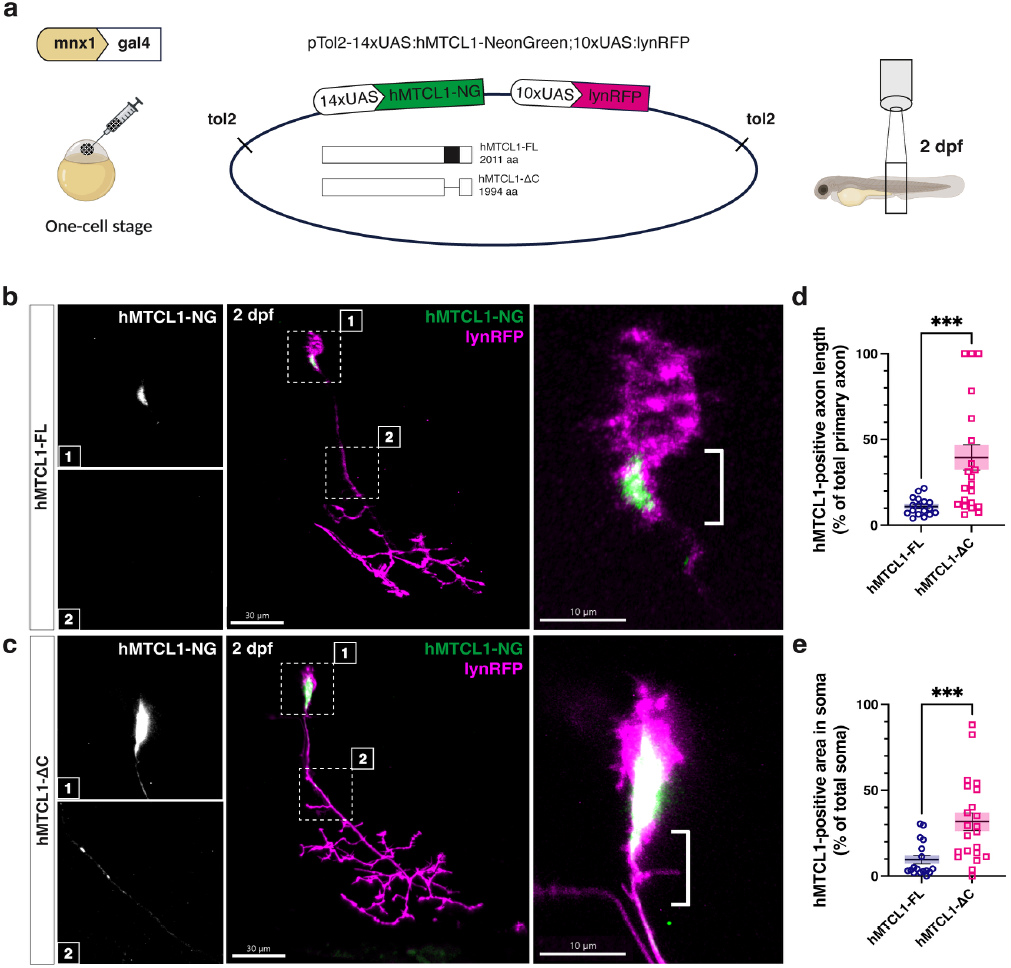
The C-terminal MTBD is required for MTCL1 localisation at the AIS. **(a)** Schematic workflow to visualize the intracellular localisation of hMTCL1. A DNA construct expressing hMTCL1 with (hMTCL1-FL) or without its C-terminal MTBD (hMTCL1-*Δ*C), fused to a fluorescent protein (NG, *NeonGreen*) and combined with a membrane reporter (*lynRFP*) under a UAS promoter, was microinjected in single-cell eggs. Single primary motor neurons could be labelled thanks to the *Tg(mnx1:gal4)* background. We imaged the zebrafish spinal cord at 2 dpf using confocal microscopy. **(b, c)** Confocal images at 2 dpf of single-labelled CaP motoneurons expressing hMTCL1-FL (b) or hMTCL1-*ΔC* (c) showing hMTCL1 signal (hMTCL1-NG, in green) in CaP motoneuron (*lynRFP*, in magenta). Scale bar = 30 mm. Higher magnifications of the hMTCL1 signal in the soma (1) and the axon (2), and of merged images with brackets around the AIS, are shown on the left and right of the panel, respectively. **(d, e)** Quantification of the hMTCL1 fluorescent signal showing that the absence of C-MTBD leads to significant mislocalisation of hMTCL1 in the axon primary branch (d, in % of total primary axon branch length, *p = 0*.*0005*) and soma (e, in % of soma area, *p = 0*.*0006*) of CaP motoneurons. Data presented as mean ± SEM (hMTCL1-FL, n = 17, hMTCL1-*Δ*C, n = 22). Statistical significance was determined by a Mann-Whitney U-test. \*\*\**p ≤ 0*.*001*.

### Loss of MTCL1^C-MTBD^ disrupts AIS integrity and axonal architecture in developing motoneurons

To examine the role of endogenous *mtcl1*^C-MTBD^ in the organism, we generated a zebrafish line carrying a constitutive inframe deletion of this domain using CRISPR-Cas9. Prior to genome editing, we confirmed *mtcl1* expression in the brain and spinal cord by hybridization chain reaction (HCR) (Supplementary Figure 10a-c, see Methods). Homozygous mutants were viable and fertile and did not display overt morphological abnormalities.

We next examined AIS integrity in developing CaP neurons using neurofascin immunolabeling as a marker of the AIS^44,45^. At 1 dpf, *mtcl1* CRISPR-ΔC mutants exhibited a markedly expanded neurofascin-positive domain, spanning (16.13 ± 1.06)% of the axon length, compared to (7.72 ± 0.53)% in WT siblings (Figure 5a, b). These results indicate a loss of proper AIS boundary definition upon deletion of the C-terminal MTBD.

**Figure 5:**
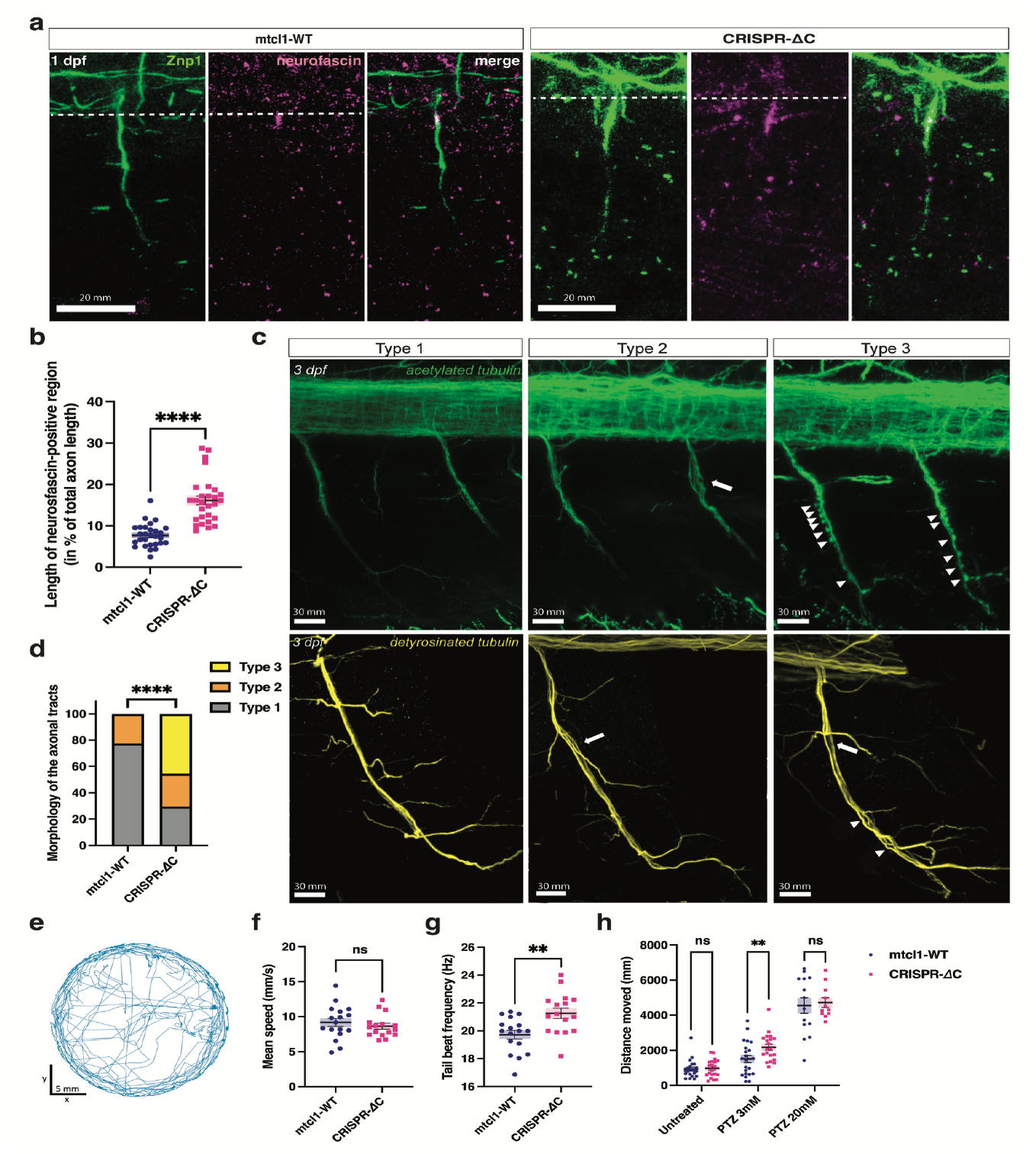
Loss of MTCL1^C-MTBD^ disrupts AIS integrity and axonal architecture in developing motoneurons, and increases seizure susceptibility. **(a)** Developing CaP neurons labelled with Znp1 (green) from control (*mtcl1-WT*, left) and mutant (*CRISPR-*Δ*C*, right) zebrafish were immunomarked for neurofascin (in magenta) at 1 dpf. The spinal cord is delimited by a white dotted line. Scale bar = 20 mm. **(b)** Quantification of the immunofluorescent signal showing significantly more spread out neurofascin along the primary axon branch in CaP neurons of *mtcl1 CRISPR-*Δ*C* mutants compared to their WT siblings (in % of total axon branch length, *p = 0*.*0001*). Data presented as mean ± SEM (*mtcl1-WT*, n = 28, *CRISPR-*Δ*C*, n = 28). Statistical significance was determined by a Mann-Whitney U-test. \*\*\*\**p ≤ 0*.*0001*. **(c)** Representative images of the three observed types of CaP motoneuron axonal tract morphologies (labelled with detyrosinated tubulin in yellow and acetylated tubulin in green). Type 1 is from control fish and types 2 and 3 are from *mtcl1 CRISPR-*Δ*C* mutant fish. White arrows point to defasciculated axons and arrow heads to axonal swellings. Scale bar = 30 mm. **(d)** Percentage of types 1, 2 and 3 axonal tract morphologies in 3 dpf control and mutant zebrafish labelled with detyrosinated tubulin (*mtcl1-WT*, n = 35; *CRISPR-*Δ*C*, n = 41; \*\*\*\**p ≤ 0*.*0001*). Data represent mean and SEM. Statistical significance was assessed by a Mann-Whitney U-test. **(e)** Exemplary 5 min-long trajectory of a freely swimming fish at 6 dpf. **(f, g)** Tracking of 6 dpf larvae freely swimming in a Petri dish showing a slightly higher tail-beat frequency (g) (*p = 0*.*0088*) in *mtcl1 CRISPR-*Δ*C* mutant fish compared to their WT siblings, despite no significant difference in mean speed (f) (*mtcl1-WT*, n = 17; *CRISPR-*Δ*C*, n = 16). Data represent mean and SEM. Statistical significance was determined by a Mann-Whitney U-test. \*\**p ≤ 0*.*01*. **(h)** Larval zebrafish locomotor behavior at 6 dpf recorded for 15 min before and after PTZ exposure to either 3 mM PTZ (*mtcl1-WT*, n = 22; *CRISPR-*Δ*C*, n = 20) or 20 mM PTZ (*mtcl1-WT*, n = 14; *CRISPR-*Δ*C*, n = 11). Locomotor behavior was quantified as the total distance moved over the recording period. Data represent mean and SEM. Statistical significance was determined by ANOVA (p=0.9028 for PTZ 20mM, ** p=0.0069 for PTZ 3mM).

Given the established role of MTCL1 in MT organization^3–5^, we analyzed axonal MT architecture using acetylated and detyrosinated tubulin labeling at 3 dpf, a time point at which no apparent morphological defects were detected. Axonal tracts were classified into distinct morphological categories (Figure 5c). In WT animals, the axons of motoneurons extend ventrally from the spinal cord and fasciculate together to form the nerve in the ventral myotome. The majority of axons in WT fish (77%) displayed an expected smooth, slender morphology (type 1), whereas this proportion was reduced to 30% in *mtcl1* CRISPR-ΔC mutants. Partial separation of the axons forming the nerve (defasciculation, type 2) was observed at similar frequencies in WT and mutant animals (23% and 25%, respectively). In contrast, axonal swellings (type 3), reminiscent of defects reported upon MTCL1 depletion in mammalian cells^5^, were observed exclusively in mutants and accounted for approximately 50% of axonal tracts (Figure 5d).

Together, these findings establish that the MTCL1^C-MTBD^ is essential for maintaining AIS integrity and axonal MT organization *in vivo*. Consistent with our structural data showing that this domain stabilizes MTs, its loss leads to defective neurofascin confinement and pronounced alterations in axonal architecture, supporting a model in which MTCL1^C-MTBD^ reinforces the specialized MT network required for AIS structure and function.

### MTCL1^C-MTBD^ deletion increases seizure susceptibility in zebrafish larvae

Given that AIS integrity and axonal MT organization are critical for proper neuronal excitability and circuit output, we next asked whether their disruption in *mtcl1* CRISPR-ΔC mutants leads to functional deficits at the behavioral level. To this end, we analyzed locomotor behavior in freely swimming larvae at 6 dpf (Figure 5e). Mutant animals displayed average and maximal swimming speeds comparable to WT siblings, but exhibited a modest yet significant increase in tail beat frequency (Figure 5f, g). This change was not associated with alterations in tail angle, bout duration, or interbout interval (Supplementary Figure 10f–j), indicating a very specific effect on motor output.

Mutations affecting MTCL1^C-MTBD^, including truncations lacking the C-terminal region, have been identified in patients with cerebellar ataxia, in some cases accompanied by seizures^5,7,46,47^. To assess whether the zebrafish CRISPR-ΔC model recapitulates this aspect of the disease, we challenged larvae with the pro-convulsant agent pentylenetetrazol (PTZ), a GABA receptor antagonist commonly used to induce seizure-like activity^48^. At a standard seizure-inducing concentration (20 mM), WT and mutant larvae exhibited comparable responses (mean distance travelled: 4556 mm vs. 4723 mm, respectively; Figure 5h and Supplementary Figure 10k). In contrast, at a sub-threshold concentration (3 mM), CRISPR-ΔC mutants displayed significantly increased swimming activity compared to WT siblings (2179 mm vs. 1504 mm over 15 minutes; Figure 5h and Supplementary Figure 10l), indicative of heightened seizure susceptibility. Thus, loss of the MTCL1^C-MTBD^ sensitizes neuronal circuits to hyperexcitability. This phenotype mirrors the seizure susceptibility reported in patients carrying MTCL1 mutations, supporting a direct link between MTCL1^C-MTBD^ dysfunction and neurological impairment. Together, these results show that loss of *mtcl1*^C-MTBD^ leads to subtle and specific changes in motor output, along with increased sensitivity to pro-convulsant stimuli, indicating that disruption of *mtcl1*-dependent MT organization at the level of the AIS predisposes motoneuron circuits to hyperexcitability in zebrafish.

## Discussion

### MTCL1 couples MT stabilization to α-tubulin acetylation and regulates neuronal physiology

Acetylation of α-tubulin at K40 is the only known PTM associated with long-lived and strain-resistant MTs^17,18^, yet how this luminal modification becomes selectively enriched on specific MT populations remains poorly understood. The causal relationship between K40 acetylation and MT stability has long been debated, with evidence suggesting that acetylation often accumulates on long-lived MTs rather than acting as a primary determinant of MT stability^10,49^. Indeed, loss of αTAT1 can increase MT stability in some contexts^50^, indicating that enrichment of acetylation may reflect local regulation of the modifying enzyme rather than a direct stabilizing effect. These observations raise the possibility that mechanisms coupling MT stabilization to local control of αTAT activity are required to generate selectively acetylated MT populations in cells.

The findings presented here identify MTCL1 as a direct regulator of this process, linking the presence of a MAP to the writing of a PTM through structural remodeling of the MT lumen. While this protein is normally enriched at the AIS, deletion of the C-MTBD causes a pronounced redistribution of the protein along the axon (Figure 4), indicating that this domain is required for proper subcellular targeting of MTCL1 within neuronal MT arrays. The structural interface identified in our cryo-EM analysis not only mediates engagement with the MT lattice *in vitro* but also appears essential for positioning MTCL1 within specialized MT architectures *in vivo*.

Consistent with this observation, zebrafish carrying an endogenous deletion of the C-MTBD exhibit early alterations in AIS organization, evidenced by a broadened distribution of an AIS marker in neurons (Figure 5a). AIS identity depends on the organization of stable MT populations, suggesting that MTCL1 contributes to maintaining the cytoskeletal architecture that defines this compartment.

At the molecular level, our results show that MTCL1 alters the accessibility of the α-tubulin H1-S2 loop to acetyltransferases, through a mechanism involving molecular mimicry between MTCL1 and α-tubulin. This internal loop, which normally constrains recognition and catalysis due to its limited conformational freedom^16^, becomes more available to the “PTM-writer” enzyme by sampling a broader ensemble of states (Figure 3f-h). *In vitro* and *in cellulo* experiments from this work show that MTCL1 enhances acetylation, while mutations that disrupt luminal remodeling attenuate the effect (Figure 3a-e). Together, these results define a structural mechanism for MAP-mediated regulation of a luminal PTM and reveal neuronal phenotypes consistent with broader physiological consequences, although a direct link remains to be established. As development proceeds, mutant axons display spheroid-like protrusions along the axonal shaft that are apparent upon staining for acetylated, but not detyrosinated, MTs (Figure 5c, Supplementary Figure 10d), indicating a selective perturbation of the acetylated MT population. The absence of the synaptic vesicle marker SV2 from these spheroids (Supplementary Figure 10d) argues against ectopic synapse formation and instead suggests the accumulation of cytoplasmic components, potentially related with impaired axonal transport and disorganization of MT bundles. Similar axonal abnormalities have been reported upon disruption of MTCL1 function in mammalian neurons, supporting a conserved role in maintaining axonal MT integrity^5^.

In mice, MTCL1 is highly expressed in cerebellar Purkinje cells, where its C-MTBD is required for proper localization of the AIS organizer ankyrin G and for the maintenance of bundled MT arrays within the AIS; loss of MTCL1 leads to disorganization of AIS MT bundles and locomotor defects^5^. Genetic studies further link MTCL1 dysfunction to human disease. Sequencing of patients with spinocerebellar ataxia has identified mutations within the MTCL1 C-MTBD^5^, and additional variants include frame-shifting mutations that truncate the protein and remove the entire C-terminal region while preserving the N-terminal MT-binding domain^7^. The structural analysis presented here shows that a disease-associated region lies within the ordered MT-binding interface that stabilizes the MT lattice, suggesting that impaired MT engagement and altered regulation of MT acetylation contribute to the cellular pathology observed in these conditions.

Interestingly, comparable phenotypes have also been observed in genetic models affecting seemingly unrelated pathways. In particular, knockout of the neuronal RNA-binding protein Elavl3 (HuC) induces alternative splicing of AnkG and shortened AIS structures, accumulation of axonal spheroids in Purkinje cell axons and produces cerebellar degeneration^51^. The convergence of these phenotypes suggests that neuronal polarity and axonal integrity depend on coordinated regulation of both MT stability and local post-transcriptional programs. Within this broader framework, MTCL1 may act on the structural side of this axis by organizing long-lived MT arrays and promoting their acetylation, thereby supporting the persistence and mechanical resilience of axonal MT tracks. Consistent with this view, *mtcl1*^C-MTBD^ mutant zebrafish also display subtle behavioral changes and increased sensitivity to the convulsant drug PTZ, pointing to altered motoneuron excitability or connectivity downstream of the AIS and potential defects in axonal cytoskeletal organization. These observations connect perturbation of MTCL1-dependent MT organization to functional consequences at the organismal level.

### Copolymerization programs MT lattice state and chemical identity

Strikingly, the molecular remodeling of the MT lumen does not appear to be an intrinsic property of MTCL1 binding per se, but depends on when the protein engages the polymer. Our data indicate that MTCL1 interacts with MTs through distinct binding modes that are determined by the timing of association. When binding to pre-assembled MTs, MTCL1 behaves as a classical stabilizing MAP that reinforces lateral tubulin contacts. In contrast, when present during MT polymerization, MTCL1 adopts an extended luminal binding footprint on the MT, displaying molecular mimicry with α-tubulin without disrupting overall MT lattice organization.

These observations suggest that copolymerization can function as a structural regulatory switch that programs MT lattice properties at the time of assembly *in vitro*; whether neurons deploy this mechanism remains to be determined. Most structural analyses of MAP–MT interactions rely on post-assembly binding, implicitly assuming that binding mode is independent of assembly history. However, earlier biochemical studies have hinted at assembly-dependent MAP behavior. Tau, for example, was shown to adopt two distinct binding modes: a dynamic interaction with the surface of pre-assembled MTs and a more stable association when tau is present during MT polymerization^36^. Yet, direct structural visualization of assembly-dependent MAP binding at high resolution has remained largely unexplored. More broadly, several MAPs are known to preferentially recognize specific structural states of the MT polymerization cycle. Doublecortin (DCX), for instance, binds polymerization intermediates at growing MT ends and recognizes the curved geometry characteristic of assembling 13-protofilament MTs^52^. Similarly, EB proteins selectively track MT plus ends by sensing the nucleotide state of the lattice and preferentially interacting with the GTP-rich region near the growing tip rather than with the mature GDP lattice^53^. The findings presented here extend this principle by showing that co-assembly with MTCL1 generates a structurally distinct lattice configuration that includes remodeling of the MT lumen. In this context, assembly history directly influences MT chemical identity by modulating accessibility of a luminal PTM site.

Such a mechanism is particularly relevant in neurons, where long-lived, non-centrosomal MT arrays must be both structurally stable and resilient to mechanical stress. The AIS, in particular, contains highly organized MT bundles that function as both a structural boundary and a platform for polarized transport^54^. Our study sheds light on how these MT populations acquire both stability and acetylation marks, yet the precise link between the described molecular mechanisms and the *in vivo* phenotypes needs to be further explored.

MTCL1 lowers the critical concentration for tubulin polymerization and promotes de novo MT assembly (Supplementary Figure 1b), conditions that could favor early engagement during MT growth and populate the MTCL1 binding mode that enhances acetylation. MTCL1 is enriched in cellular systems dominated by persistent MT arrays that show elevated acetylation levels^3–5^, raising the possibility of a general MTCL1-driven mechanism that coordinates the formation of long-lived MT bundles with the establishment of their chemical identity across different cell types.

## Conclusion

Our work provides a novel mechanistic framework for the long-standing observation that stabilized MT populations are frequently enriched in acetylation. In the context of MTCL1, this protein contributes to MT longevity not only by reinforcing lattice stability, but also by promoting an α-tubulin conformational ensemble that favors the deposition of a mechanically protective PTM. By engaging MTs during polymerization, MTCL1 programs a lattice state that is both structurally stabilized and permissive for α-tubulin acetylation. In this view, MTCL1 acts as a licensing factor that links MT assembly history, lattice architecture, and the spatially restricted activity of a tubulin-modifying enzyme. By coordinating MT structure, mechanics, and chemical modification across scales, MTCL1 emerges as a key organizer of long-lived, acetylation-rich MT arrays and illustrates how “writing” of the tubulin code can be modulated by MAPs in a spatially and temporally controlled manner.

This framework integrates multiple levels of biological organization, from molecular-resolution structural insights to functional consequences in a living organism. Structural observations informed a unifying model that was subsequently tested across biochemical, cellular, and *in vivo* systems. Perturbation of MTCL1 function in zebrafish revealed a clear physiological sensitivity to disruption of this pathway, linking defined structural features of the MT lattice to emergent properties of MT organization and neuronal function *in vivo*.

## Acknowledgements

We thank Z. Yang for helpful discussions. We are grateful to D. Toso and R. Thakkar from the Cal-Cryo EM facility, and to P. Tobias and K. Stine for computational support. We also thank all members of the Del Bene lab for fruitful discussions, particularly Elena Putti and Matthieu Tuffery for the behavioral setup. We acknowledge the Institut de la Vision for animal and imaging facilities, especially Stephane Fouquet and Nermine Saidi for their constant help and advice in confocal imaging. We thank Matthew Rasband for providing the anti-FIGQY neurofascin antibody. We also acknowledge the UCSF ChimeraX team for the software used in structural rendering. The acetyltransferase schematic used in the acetylation assay figure was sourced from NIAID Visual & Medical Arts (10/7/2024), IRF3, NIAID NIH BIOART Source.

This work was funded by the European Research Council (ERC-2022-SYG) under the European Union’s Horizon 2020 research and innovation programme (grant agreement N°101071583, “TUBULINCODE,” to E.N., Z.L., C.J., and F.D.B.), and by the HORIZON-MSCA-2024-PF-01 fellowship N°101207124 (“AxTransCode” to D.L.). Additional support was provided by IHU FOReSIGHT (ANR-18-IAHU-0001) (F.D.B.), and by CNRS, INSERM, and Sorbonne Université core funding (F.D.B.). T.M.J.D. was supported by FRM PhD funding ECO202206015510 and fourth-year funding FDT202504020880. Z.L. acknowledges institutional support from CAS (RVO: 86652036), the Imaging Methods Core Facility at BIOCEV supported by MEYS CR [LM2023050, Czech-BioImaging], and CF Protein Production of CIISB, Instruct-CZ Centre, supported by MEYS CR [LM2023042] and the European Regional Development Fund project “UP CIISB” [No. Z.02.1.01/0.0/0.0/18_046/0015974]. This research was supported in part by the University of Pittsburgh Center for Research Computing and Data, RRID:SCR_022735, through the resources provided. Specifically, this work used the H2P cluster, which is supported by NSF award number OAC-2117681 (to M. Gu.). Additional support was provided by start-up funding from the University of Pittsburgh School of Medicine (to M. Gu.). Further funding was provided by the National Natural Science Foundation of China (NSFC Young Student Basic Research Program, grant 224B2707 to J.L.) and the Research Grants Council of Hong Kong (Collaborative Research Fund C7064-22GF to S.C.T., as project coordinator). E.N. is a Howard Hughes Medical Institute Investigator.

## Author contributions

J.M.P.B. and E.N. initially conceived the project and all authors contributed to its development and the design of experiments. J.M.P.B. purified proteins, performed cryo-EM imaging and data processing, *in vitro* acetylation assays, molecular modeling and analyzed data. T.M.J.D. performed zebrafish genetic manipulation, *in vivo* imaging and behavioral experiments with the associated data-analysis. M.Go. and M. Gu. performed MD simulations and analyzed data. E.L. purified protein, performed TIRF dynamic MT imaging and analyzed data. D.L. performed *in cellulo* acetylation assays and analyzed data. V.H. performed molecular cloning. J.L. purified proteins. R.Z. assisted with cryo-EM data processing. E.N., F.D.B., C.J., M.Gu., S.C.T. and Z.L. secured funding. C.J., E.N., F.D.B., M. Gu. and Z.L. supervised different aspects of the project. E.L., D.L., T.M.J.D., J.M.P.B., M.Go. and M. Gu., initially wrote the manuscript, with further edits by all authors.

## Materials and Methods

### Protein expression and purification

The C-terminal MT-binding domain of human MTCL1 (residues 1678– 1773, MTCL1^C-MTBD^) was cloned from human cDNA into a pRSFDuet-SUMO vector to generate an N-terminal His-SUMO fusion. Cloning was performed by sequence and ligase independent cloning (SLIC), and the construct was validated by full plasmid sequencing (service provided by Plasmidsaurus).

For protein expression, the plasmid was transformed into Rosetta2 (DE3) competent cells, plated for kanamycin selection, and a single colony was grown in LB medium. Cultures were cooled down at OD_600_ ∼0.6, induced with 0.2 mM IPTG, and incubated at 18 °C overnight (∼18 h) before harvest. Cell pellets were lysed by sonication in His lysis buffer (20 mM HEPES pH 7.5, 300 mM KCl, 1 mM MgCl_2_, 5% glycerol, 20 mM imidazole, 0.1% Tween-20, 10 mM β-mercaptoethanol, 1× benzonase, and 1 EDTA-free protease inhibitor tablet) and clarified by centrifugation at 18,000 rcf for 1 h at 4 °C. The supernatant was incubated with Ni-NTA agarose resin equilibrated in the same buffer, washed extensively with alternating high-salt (700 mM NaCl) and regular wash buffer, and eluted by overnight on-column cleavage with Ulp1 protease to remove the His-SUMO tag. The eluate was further incubated with fresh Ni-NTA resin to deplete any uncleaved fusion protein and free SUMO.

The resulting flow-through was diluted fourfold with ion-exchange buffer A (20 mM HEPES pH 7.5, 100 mM NaCl, 10 mM β-mercaptoethanol) and loaded onto a HiTrap SP HP column. Protein was eluted with a linear salt gradient up to 1 M NaCl, and MTCL1-containing fractions were pooled, concentrated, and buffer-exchanged into SEC buffer (20 mM HEPES pH 7.5, 300 mM KCl, 1 mM MgCl_2_, 5% glycerol, 1 mM TCEP). Final purification was performed by size-exclusion chromatography on a Superdex 75 Increase 10/300 GL column equilibrated in SEC buffer. Peak fractions were concentrated, aliquoted, and flash frozen in liquid nitrogen for storage at – 80 °C.

For the dynamic MT experiments, Neon Green-labelled MTCL1^C-MTBD^ (human MTCL1 1678-1773; NM_001378205.1) was expressed in E. coli BL21(DE3)-RIPL strain. The cells were grown at 30°C until OD_600_ of 0.5-0.6, the protein expression was then induced by 0.1 mM IPTG, and the cells were grown overnight at 16°C. Bacterial cells were lysed in 20 ml of lysis buffer (50 mM Hepes pH 7.4, 300 mM KCl, 2mM MgCl_2_, 0.1% Tween, 5% glycerol, 1mM DTT, 0.1mM ATP, 1x Protease inhibitor cocktail and 1.25 µL Benzonase), sonicated on ice and centrifuged (40000 x g; 30 min; 4 °C).

The soluble fraction was incubated with HisTrap Ni-NTA agarose resin (XF340049, Thermo Scientific) for two hours at 4 °C with slow rotation. After incubation, the beads were washed first with 20 ml of Wash Buffer I (50 mM HEPES, pH 7.4; 300 mM KCl; 0.1% Tween; 5% glycerol; 1 mM DTT; 0.1 mM ATP; 30 mM imidazole), and then with 10 ml of Wash Buffer II (Wash Buffer I with 60 mM imidazole). The His-trapped proteins were then eluted with elution buffer (wash buffer I containing 250 mM imidazole). The eluted fractions were subjected to Strep-Tactin XT purification (washing buffer: 50 mM Hepes pH 7.4, 300 mM KCl, 2 mM MgCl_2_, 1 mM EGTA, 0.1% Tween, 5% glycerol, 1 mM DTT and 0.1 mM ATP) and eluted using elution buffer (washing buffer with 50 mM biotin). The eluted proteins were then dialyzed overnight using dialysis buffer (50 mM Hepes pH 7.4, 300 mM KCl, 0.1% Tween, 5% glycerol, 30 mM imidazole) in dialysis tubes with a 12 kDa cut-off, in the presence of C3 protease (for cleavage of the 6xHis-tag and Strep-tag). After dialysis, the solution was loaded onto Ni-NTA agarose resin, and the flow-through was collected to remove the C3 protease (6xHis-tagged). Protein concentration was measured with a NanoDrop at 280 nm absorbance. Proteins were flash-frozen in liquid nitrogen and stored at -80°C. All purification steps were performed at 4°C. Recombinant human α1Aβ4B tubulin dimers and C. elegans αTAT2 were purified as previously described ^16^.

### Co-sedimentation assay

10 uL of 10 mg/mL porcine brain tubulin (Cytoskeleton Cat # T240) was polymerized in the presence of 1 mM GTP at 37 °C for 1 hour. Following polymerization, MTs were pelleted to remove polymerization-incompetent tubulin and subsequently resuspended in pre-warmed cryo buffer (BRB80 supplemented with 0.05% NP-40, 1.5 mM MgCl_2_, 1 mM DTT). The MT concentration was measured and diluted to 2 µM, a concentration below the critical threshold for spontaneous polymerization^55,56^, at which MTs are expected to depolymerize even at 37 °C. Diluted MTs were incubated at 37 °C with different ratios of MTCL1^C-MTBD^, which had been desalted to cryo buffer using a Zeba desalting column. Samples were then subjected to ultracentrifugation at 45,000 rpm for 40 minutes at 37 °C. No Taxol or any other MT-stabilizing agents were used in these experiments.

### Cryo-EM sample preparation

For pre-assembled MTs, porcine brain tubulin (Cytoskeleton Cat # T240) was reconstituted to 10 mg/mL in BRB80 buffer (80 mM Pipes pH 6.9, 1 mM EGTA, 1 mM MgCl_2_) supplemented with 10% glycerol, 1 mM GTP, and 1 mM DTT. Tubulin was polymerized at 37 °C for 30 min in the presence of 2 µL of 2 mM Taxol. MTs were pelleted by centrifugation at 37 °C and 15,000 rcf for 20 min, the supernatant was discarded, and the pellet was resuspended in resuspension buffer (BRB80 supplemented with 0.05% NP-40, 1.5 mM MgCl_2_, 1 mM DTT, 250 µM Taxol). Tubulin concentration was measured in a CaCl_2_-depolymerized aliquot and the MTs were diluted to 2 µM in dilution buffer (BRB80 supplemented with 0.05% NP-40, 1.5 mM MgCl_2_, 1 mM DTT, 100 µM Taxol). Immediately prior to sample preparation, MTCL1^C-MTBD^ was desalted into cryo buffer using Zeba spin columns. For grid preparation, 2 µL of 2 µM Taxol-stabilized MTs were applied to glow-discharged holey carbon grids (QuantiFoil, Cu 300 R 2/1), incubated for 30 s, manually blotted, and incubated with 2.5 µL of 190 µM MTCL1^C-MTBD^. Grids were transferred to a Vitrobot (Thermo Fisher Scientific) set at 25 °C and 80% humidity, incubated for 1 min, blotted (blot force 6 pN, blot time 6 s), plunge-frozen in liquid ethane, and stored in liquid nitrogen.

For sample preparation under copolymerization conditions, an equimolar mixture of MTCL1^C-MTBD^ and porcine brain tubulin (12 µL total) was centrifuged at 100,000 rcf for 15 min at 4 °C to remove aggregates. The supernatant was transferred to a fresh tube, and in the case of the +Taxol condition, 2 µL of 2 mM Taxol were added during polymerization at 37 °C for 30 min. Samples were then centrifuged at 100,000 rcf for 20 min at 30 °C to pellet MTs, and resuspended in 40 µL of resuspension buffer (BRB80 supplemented with 0.05% NP-40, 1.5 mM MgCl_2_, 1 mM DTT, 250 µM Taxol). From this point, sample preparation on grids was carried out as described above for the pre-assembled MT dataset.

### Cryo-EM sample preparation

For pre-assembled MTs, porcine brain tubulin (Cytoskeleton Cat # T240) was reconstituted to 10 mg/mL in BRB80 buffer (80 mM Pipes pH 6.9, 1 mM EGTA, 1 mM MgCl_2_) supplemented with 10% glycerol, 1 mM GTP, and 1 mM DTT. Tubulin was polymerized at 37 °C for 30 min in the presence of 2 µL of 2 mM Taxol. MTs were pelleted by centrifugation at 37 °C and 15,000 rcf for 20 min, the supernatant was discarded, and the pellet was resuspended in resuspension buffer (BRB80 supplemented with 0.05% NP-40, 1.5 mM MgCl_2_, 1 mM DTT, 250 µM Taxol). Tubulin concentration was measured in a CaCl_2_-depolymerized aliquot and the MTs were diluted to 2 µM in dilution buffer (BRB80 supplemented with 0.05% NP-40, 1.5 mM MgCl_2_, 1 mM DTT, 100 µM Taxol). Immediately prior to sample preparation, MTCL1^C-MTBD^ was desalted into cryo buffer using Zeba spin columns. For grid preparation, 2 µL of 2 µM Taxol-stabilized MTs were applied to glow-discharged holey carbon grids (QuantiFoil, Cu 300 R 2/1), incubated for 30 s, manually blotted, and incubated with 2.5 µL of 190 µM MTCL1^C-MTBD^. Grids were transferred to a Vitrobot (Thermo Fisher Scientific) set at 25 °C and 80% humidity, incubated for 1 min, blotted (blot force 6 pN, blot time 6 s), plunge-frozen in liquid ethane, and stored in liquid nitrogen.

For sample preparation under copolymerization conditions, an equimolar mixture of MTCL1^C-MTBD^ and porcine brain tubulin (12 µL total) was centrifuged at 100,000 rcf for 15 min at 4 °C to remove aggregates. The supernatant was transferred to a fresh tube, and in the case of the +Taxol condition, 2 µL of 2 mM Taxol were added during polymerization at 37 °C for 30 min. Samples were then centrifuged at 100,000 rcf for 20 min at 30 °C to pellet MTs, and resuspended in 40 µL of resuspension buffer (BRB80 supplemented with 0.05% NP-40, 1.5 mM MgCl_2_, 1 mM DTT, 250 µM Taxol). From this point, sample preparation on grids was carried out as described above for the pre-assembled MT dataset.

### Cryo-EM data collection

Data for MTs decorated with MTCL1C-MTBD were collected using an Arctica (pre-assembled Taxol MTs + MTCL1C-MTBD and copolymerized MTs + MTCL1C-MTBD) or Titan Krios (copolymerized Taxol MTs + MTCL1C-MTBD) microscopes (Thermo Fisher Scientific) operated at 200 or 300 kV, respectively (Supplementary Table 1). All cryo-EM images were acquired on a K3 direct electron detector (Gatan) at nominal magnifications of 36,000× (Arctica) and 81,000x (Titan Krios), corresponding to calibrated physical pixel sizes of 1.14 and 1.05 Å, respectively. Depending on the dataset, the detector was operated either in super resolution or regular counting mode. The dose was fractionated into 50 frames, with a total accumulated dose of ∼50 electrons/Å^2^ on the specimen. All data were collected semi-automatically using the SerialEM software package^57^.

### Cryo-EM image processing

For the MTCL1^C-MTBD^ datasets, movie stacks were imported into Relion 5^58^ and motion-corrected. The motion-corrected micrographs were moved into CryoSparc for further processing. CTF parameters were estimated with Patch CTF, and micrographs were manually curated to remove poor-quality images. Segments were automatically picked with the filament tracer — initially reference-free, followed by template-based picking using selected 2D classes. The inter-pick distance was set to 82 Å, corresponding to one αβ-tubulin dimer. Particles were extracted in 512-pixel boxes and Fourier-cropped to 256 pixels for initial processing. Two rounds of 2D classification were used to select classes showing clear MT features and to discard blurry or off-center segments and junk. MTs with different protofilament numbers were separated by heterogeneous refinement using low-pass filtered 12-, 13-, 14-, and 15-PF references as initial models; the majority PF species (14-PF) was retained for subsequent steps.

Selected particles were helically refined using initial parameters of 82.5 Å rise and 0° twist, followed by local refinement with a hollow cylindrical mask around the MT. To account for the symmetry-breaking seam, a Frealign-based seam-search routine was applied on a per-particle basis: CryoSPARC alignment parameters from the last local refinement were converted to STAR format with csparc2star (PyEM) and then to PAR format for Frealign using an in-house script. After seam assignment, particles were re-imported into CryoSPARC with seam-corrected orientations and re-extracted without Fourier cropping (512-pixel boxes), using the improved alignments for recentering. A local refinement was performed to obtain a C1 reconstruction, followed by local CTF refinement to estimate per-particle CTF parameters. The symmetry-search job was used to determine the refined rise and twist, which were then applied in symmetry expansion to exploit the pseudo-symmetry of each MT particle.

After symmetry expansion, refinements focused on a shaped mask encompassing four tubulin dimers — two longitudinally adjacent and two laterally adjacent — to capture the inter-dimer interfaces relevant for MTCL1 binding. The resulting consensus maps were subjected to alignment-free 3D classification to resolve compositional and conformational heterogeneity. 3D classification was implemented in a way that initial volumes were reconstructed from random particle subsets, and then iteratively improved during class assignment. This sorting procedure yielded reconstructions of MTCL1-bound MTs with short and extended footprints, as well as an empty MT class. MTCL1-containing volumes were post-processed with DeepEMhancer^59^ to bring out high-resolution features for improved visualization. Cryo-EM processing parameters are summarized in Supplementary Table 1.

### Model building and refinement

The final cryo-EM maps from both datasets —pre-assembled Taxol-stabilized MTs and copolymerized MTs prepared in the absence of Taxol— were used for model building and coordinate refinement. Four αβ-tubulin dimers from PDB 6DPV were rigid-body fitted in ChimeraX, and MTCL1^C-MTBD^ was built in Coot by manual backbone tracing and residue assignment guided by the high-resolution side-chain features present in the MTCL1 density within the inter-protofilament groove, which enabled an unambiguous sequence register. For the copolymerized structure, the H1-S2 loop in some of the α-tubulin subunits had to be partially deleted to accommodate for the extra MTCL1 density. Separate real-space refinements in Phenix were performed against each map using a model containing four αβ-tubulin dimers and two MTCL1^C-MTBD^, and refinement statistics are reported in Supplementary Table 1. For figure preparation, refined coordinates and maps were displayed in ChimeraX and colored by subunit type only, with α-tubulin in green, β-tubulin in blue, and MTCL1 in red. Lattice compaction was quantified from the refined coordinates by averaging Cα–Cα distances across corresponding α–β residue pairs that report on intra- and inter-dimer spacing, and the MTCL1-bound state was compared to the reported compaction of a physiological GDP lattice.

### MT assembly for *in vitro* TIRF assays

Porcine brain tubulin was isolated using the high-molarity PIPES procedure as described previously^60^. Biotin-labeled tubulin was purchased from Cytoskeleton Inc. (T333P). GMPCPP-MT seeds were polymerized from 4 mg/ml tubulin (1:50 biotin-labeled tubulin: unlabeled tubulin) for 4 h at 37 °C in BRB80 (80 mM PIPES, 1 mM EGTA, 1 mM MgCl_2_, pH 6.9) supplemented with 1mM MgCl_2_ and 1mM GMPCPP (Jena Bioscience, NU-405). The polymerized MTs were centrifuged for 30 min at 18000 x g in a Microfuge 18 Centrifuge (Beckman Coulter). After centrifugation the pellet was resuspended and kept in BRB80 at room temperature. Before the experiments the GMPCPP stabilized MTs were broken into short MT seeds using a Hamilton pipette.

### TIRF microscopy

TIRF microscopy was performed using an inverted microscope (Nikon Ti-E) with an H-TIRF module. Imaging was performed using 60× oil-immersion objectives with a numerical aperture of 1.49 (Apo TIRF; Nikon). Image acquisition was performed using sCMOS cameras (Hamamatsu Photonics). MTs were detected using IRM, while fluorescently labelled proteins were imaged using filter cubes corresponding to eGFP and 488 nm laser. The microscope was operated and images were acquired using NIS-Elements AR (v. 5.20).

TIRF flow chambers were prepared using thin parafilm strips and two cleaned, HMDS-treated glass coverslips (22 × 22 mm^2^ and 18 × 18 mm^2^; Corning). Chambers were incubated with anti-biotin antibody (20 µg/ml; Sigma Aldrich, B3640) in PBS for 5 min and blocked with 1% Pluronic F127 (Sigma Aldrich, P2443) for ≥30 min. After that the chambers were washed with 40uL of BRB80.

### TIRF assays

*In vitro* MT dynamics assays were performed using TIRF microscopy as described above. GMPCPP-stabilized MT seeds were first introduced into the imaging chambers. Unbound MTs were washed away with BRB80 and chambers were equilibrated with TIRF assay buffer (BRB80 supplemented with 10 mM dithiothreitol, 0.02 mg/ml casein, 1 mM Mg-ATP, 20 mM D-glucose, 1% Tween, 0.22 mg/ml glucose oxidase and 20 µg/ml catalase) prior to imaging. The polymerization buffer was prepared from the assay buffer by adding 0,1% methylcellulose, 2.5 mM GTP and 10 mM MgCl_2_.

MTCL1 interaction with dynamic MTs: For control conditions, chambers with MT seeds were loaded with a polymerization mixture consisting of 5 µl polymerization buffer, 4 µl tubulin (4 mg/ml), and 1 µl assay buffer. For MTCL1 conditions, 1 µl assay buffer was replaced with 1 µl MTCL1 (final concentration 600 nM). The resulting polymerization mixes were introduced into the respective chambers, and MT growth dynamics were recorded for 25 min with images acquired every 5 s. The experiment was repeated three times.

MTCL1 effect on MT behaviour after washout: Dynamic MTs were assembled as described for the MT dynamics experiment above, in the presence and absence of MTCL1. Briefly, polymerization mixtures were introduced into flow chambers and MTs were allowed to grow for 20 min. Subsequently, the chambers were washed with 30 µL of assay buffer. MT behaviour following the washout was then monitored by time-lapse imaging. Images were acquired every 5 s, and the experiment was repeated three times.

### TIRF image analysis

Microscopy data were analyzed using ImageJ (version 2.16.0/1.54q, Fiji). Kymographs of individual MTs were generated by drawing a line along the MT lattice and using the ImageJ KymographBuilder plugin, for both control and MTCL1 conditions. Kymographs were subsequently analyzed to obtain MT dynamic parameters.

For MT dynamics experiments, the following parameters were quantified: (i) growth rate (nm/s), (ii) depolymerization rate (nm/s), (iii) catastrophe frequency (s^-1^), and (iv) rescue frequency (s^-1^). For washout experiments, MT depolymerization rates (nm/s) after washout were measured. A total of 48 MTs were analyzed for control conditions and 42 for MTCL1 conditions in the MT dynamics experiments, pooled from three independent experiments. For washout experiments, 40 MTs (control) and 39 MTs (MTCL1) were analyzed, also from three independent experiments. Statistical analysis was performed using unpaired two-tailed t-tests in GraphPad Prism (version 10.6.1). Differences were considered statistically significant when p < 0.05 (*p < 0.05; **p < 0.01; ***p < 0.001; ****p < 0.0001). Data are presented as individual data points with the mean indicated.

### Acetylation assays

For acetylation assays, α1Aβ4B tubulin, MTCL1, and αTAT2 stock solutions were first clarified by ultracentrifugation at 287,000 rcf to remove aggregates formed during thawing. Protein concentrations were then determined using the Bradford assay. MT substrates were prepared by polymerizing 11 µM α1Aβ4B recombinant tubulin in the presence or absence of 40 µM MTCL1, with 1 mM GMPCPP included in all conditions (polymerization buffer: 1x BRB80, 1.5 mM MgCl2, 10% glycerol). In samples lacking MTCL1, an equivalent volume of MTCL1 buffer (20 mM HEPES pH 7.5, 300 mM KCl, 1 mM MgCl_2_, 5% glycerol, 1 mM TCEP) was added to control for differences in ionic strength and buffer composition. Polymerization reactions were incubated at 37 °C for 2 hours to ensure complete assembly. Following polymerization, MTs were pelleted by ultracentrifugation at 37 °C and 287,000 rcf for 15 minutes and resuspended in 20 µL of warm assay buffer (1x BRB80, 1 mM TCEP). The concentration of polymerized MTs was determined by Bradford assay after depolymerizing a small aliquot on ice in the presence of CaCl_2_.

Acetylation reactions were carried out using 3 µM MT substrate (with or without MTCL1), supplemented with 0.1 mg/mL free tubulin, and 100 µM acetyl-CoA lithium salt. Reactions were initiated by the addition of 0.25µM αTAT2 and incubated at 37 °C. At defined time points (0, 15, 30, 60, 120, and 240 minutes), 2 µL aliquots were withdrawn and immediately quenched in 48 µL of 1× SDS loading buffer. Acetylation levels and total tubulin content were assessed by immunoblotting against acetylated K40 α-tubulin (Thermo Fisher Scientific, 6-11B-1) and Coomassie Blue staining, respectively.

Time-course acetylation data were analyzed by fitting the normalized acetylation values to a single-exponential saturation model of the form

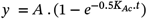

where *y* is the normalized acetylation fraction, A is an amplitude factor, *t* is time (in minutes), and K_Ac_ is the apparent acetylation constant in min^-1^.

For each condition, mean values and standard errors were calculated from at least 3 independent replicates at each time point, and the fits were performed using non-linear least-squares regression. The error in K_Ac_ was derived from the covariance matrix of the fit and reported as the standard error of the fitted parameter. Fitted curves were plotted together with the mean ± SE experimental data.

### Immunofluorescence in cells

HeLa cells were seeded onto glass coverslips in 24-well plates in DMEM (Gibco, Thermo Fisher Scientific) supplemented with 10% FBS (v/v) and Penicillin-Streptomycin (Life Technologies). The following day, the cells were transfected with GFP, GFP-FL hMTCL1, GFP-FL hMTCL1 ΔN, GFP-hMTCL1^C-MTBD^ or GFP-hMTCL1^C-MTBD^ AKR using jetPEI (according to manufacturer’s guidelines, with 0.75 µg of DNA and 1.5 µl of jetPEI in plain DMEM). 24 h after transfection, the cells were fixed for 10 minutes using 4% paraformaldehyde (PFA), 0.3% glutaraldehyde + 0.5% Triton X-100 in cytoskeleton buffer (CB) containing glucose (10 mM MES, 150 mM NaCl, 5 mM EGTA, 5 mM MgCl2, 5mM glucose, pH 6.1). After washes with PBS, the cells were treated with sodium borohydride solution (0.1% NaBH4/PBS) to reduce background fluorescence. Then, the samples were incubated with blocking solution (5% BSA in CB + 0.05% Tween-20) for 1 h at room temperature (RT), followed by incubation with the primary antibodies mouse anti-acetylated α-tubulin (acetyl K40; [6-11B-1]; 1:500; #T6793, Sigma-Aldrich) and rat anti-α-tubulin, tyrosinated (clone YL1/2; 1:3000; #MAB1864, Merck-Millipore), diluted in blocking solution for 1h RT. After washes with PBS (1× for 5min) and PBS containing 0.1% Tween-20 (PBST) (2× for 5 min), the cells were incubated with secondary antibodies Alexa Fluor 568 and 647 (1:1,000; Thermo Fisher Scientific; diluted in blocking solution) for 1 h RT, followed by final PBST and PBS washes. The cells were imaged (z-planes separated by 0.23 µm) in a ZEISS LSM 980 confocal microscope, using the 63× oil-immersion objective (N.A. 1.40), a ZEISS Axiocam 820 mono camera, 488-nm, 561-nm and 639-nm lasers and Zen software (Carl Zeiss). Images were analyzed using ImageJ software. Integrated fluorescence density was quantified from sum-projected images by drawing a freehand ROI around each cell. Background fluorescence was measured using the same ROI placed outside the cell and subtracted from the corresponding cell signal. α-tubulin acetylation levels were normalized to α-tubulin tyrosination, and the ratio normalized to the average level of FL hMTCL1 cells. Representative images were 3D LSM Plus processed using Zen software and analysed using ImageJ (maximum-intensity projection of z-stacks).

### Molecular dynamics simulations

All-atom MD simulations of αTAT2 bound to MTs were initiated from the structure with PDB ID 8Y9F, in which αTAT2 is bound to the MT and Lys40 is acetylated in a catalytically relevant conformation. Acetyl-CoA was taken from the PDB ID 8YAR structure, which was the enzymatically locked state containing the Lys40Arg mutation (set 1, three independent runs, 900 ns total). A second system (set 2, three independent runs, 2.25 µs length in total) was generated by removing αTAT2 from the PDB 8Y9F^16^ structure. For the third set of simulations (set 3, three independent runs, 2.25 µs total,), the missing tubulin region Pro37–Gly59, which contains the H1–S2 loop at the MTCL1^C–MTBD^ binding site, was modeled as follows: We took an H1–S2 loop conformation from our previous tubulin-only simulations^61^ and superimposed the corresponding tubulin dimer onto the MT lattice of PDB 5SYF3^62^. We then steered the center of mass of this loop toward the center of the MT (i.e., along the radial direction). This loop conformation was added to our MTCL1^C–MTBD^-tubulin structure and the resulting protein system was used as the starting structure for the MTCL1^C–MTBD^–bound simulations. The fourth and fifth sets of simulations (sets 4 (three independent runs, 1.5 µs length in total) and 5 (seven independent runs, 2.1 µs length in total)) were initiated by placing αTAT1 10 Å away from the tubulin surface along a vector approximately orthogonal to the αTAT1– tubulin binding interface. In set 4, the position of αTAT1 was restrained to maintain this fixed separation from the MT, whereas in set 5 αTAT1 was allowed to move freely and bind.

Each system was solvated using the TIP3P water model in a cubic box chosen to provide at least 15 Å of water between any protein atom and the box edge. K^+^ and Cl^−^ ions were added to neutralize the system and to reach a final salt concentration of 150 mM KCl. This procedure yielded total system sizes of approximately 300,000, 350,000, 400,000, 400,000, and 400,000 atoms for sets 1–5, respectively. MD simulations were performed using NAMD 3^63^ with the CHARMM36m^64^ all-atom additive protein force field. A 2-fs integration time step was used, and simulations were carried out at 310 K and 1 atm. Long-range electrostatics were treated with the particle-mesh Ewald method, and a 12 Å cutoff was applied for van der Waals interactions.

To equilibrate each system, all protein atoms were first held fixed during 10,000 steps of energy minimization, followed by 1 ns of equilibration of the solvent. The protein restraints were then released, and the system underwent an additional 10,000 steps of minimization. Subsequently, a 5-ns equilibration was performed while restraining protein Cα atoms with a harmonic potential of 1 kcal mol^-1^ Å^-2^. Following these two minimizations– equilibration phases, production simulations were initiated. To maintain the integrity of the MT lattice, Cα atoms of selected tubulin residues (Supplementary Table 2) were restrained with a harmonic potential of 1 kcal mol^-1^ Å^-2^ throughout the production runs.

### Fish lines and husbandry

Zebrafish (*Danio rerio*) were maintained under controlled conditions, at a temperature of 28°C and subjected to a 14-hour light/10-hour dark cycle. Embryos were collected and cultured in fish water supplemented with 0.003% 1-phenyl-2-thiourea to prevent pigmentation and 0.01% methylene blue to inhibit fungal growth in petri dishes. Fish were housed within our institute’s animal facility, built following local animal welfare standards. All animal procedures adhered to the ethical guidelines outlined by French and European Union regulations. Animal handling and experimental procedures were approved by the committee on ethics of animal experimentation of Sorbonne Université (APAFIS#21323-2019062416186982). Experimental procedures were conducted on larval zebrafish before the onset of sexual differentiation. The following transgenic and mutant fish lines were used: *Tg(mnx1:gal4)* (Bercier et al. 2019) and *mtcl1-*Δ*C*.

### Zebrafish mutants

The gRNAs used to generate the different CRISPR in-frame deletion are the following: gRNA-mtcl1-ΔC-1 5’-AAGGCAATAACGAATCCGCTTGG-3’ and gRNA-mtcl1-Δ*C-2* 5’-GCTGGAGAGGCCGTCGTTGATGG-3’. Primers used for genotyping are listed below: mtcl1_ex15_FW (5’-GTAAGACCAACAGAGTGAGTGG-3’) and mtcl1_ex15_RV (5’-CTTCGAAGCCTGTTGTGATTGAGG-3’). The *mtcl1-*Δ*C* mutant line was generated using the CRISPR/Cas9 gene editing technology. Two gRNAs were injected to target exon 15 of the *mtcl1* gene leading to a 48 bp deletion, causing an in-frame deletion in the C-MTBD of the endogenous zebrafish *mtcl1* (Supplementary Figure 10c). PCR amplicons were deposited on 2% agarose gel for the identification of wild type and mutants.

### Molecular cloning for *in vivo* experiments

The *ubc* intron was amplified by PCR from the *pTol1-14UAS:ubc-EGFP-myo1b* from Revenu et al. (2024)^65^. The *mNeonGreen* fragment was amplified from an in-house plasmid. Codon-optimized *hMTCL1-FL* and *hMTCL1-ΔCter* cDNA were amplified from a construct obtained from VectorBuilder (#VB240812-1510zmn). The *10UAS:lynRFP* construct used in this study was previously cloned by Auer and Del Bene (2014)^66^. Fragments were subsequently subcloned into the *pTol1-14UAS* plasmid using the Gibson Assembly Cloning Kit (New England Biolabs) following manufacturer’s instructions to generate *pTol1-14UAS:ubc-hMTCL1-FL-mNeonGreen;10UAS:lynRFP* and *pTol1-14UAS:ubc-hMTCL1-ΔCter-mNeonGreen;10UAS:lynRFP*.

### Micro-injections

Fertilized eggs from the *Tg(mnx1:gal4)* fish line were collected in egg water and aligned into prepared 1.5% agarose injection molds (agarose diluted in water). To inject, a pressure injector (FemtoJet microinjector, Eppendorf) with borosilicate glass capillaries (GC100TF10, Clark Electromedical Instruments) was used. The capillaries were previously filled with the following injection solutions and injected through the chorion into the single cell cytoplasm. The *pTol1-14UAS:ubc-hMTCL1-FL-mNeonGreen;10UAS:lynRFP* and *pTol1-14UAS:ubc-hMTCL1-ΔCter-mNeonGreen;10UAS:lynRFP* plasmids were injected at 25 ng/µl with 25 ng/µl of *tol1* mRNA, for sparse cellular labelling.

CaP neurons were selectively labeled under the *mnx1* promoter, together with a membrane-targeted fluorescent reporter (10UAS*:lyn*RFP) to confirm neuronal identity based on morphology^67^. Expression of the hMTCL1 constructs was driven by a 14×UAS promoter following single-cell stage injection, with each construct fused to a *NeonGreen* fluorescent tag (Figure 4a).

### Immunohistochemistry

Embryos were fixed at 1 dpf and 3 dpf in freshly made 4% paraformaldehyde diluted in PBS for 2h at room temperature. They were then rinsed multiple times in 1X PBS containing 0.1% triton X-100 (PBS-t).

Immunohistochemistry experiments were performed as follows: whole larvae were washed three times in PBS-t solution. Samples were permeabilized using proteinase K (Sigma, 20ug/ul) for 8 to 70 min according to their developmental stage and postfixed in 4% paraformaldehyde diluted in PBS for 20 min. Consecutively, samples were rinsed in PBS-t 2 times and incubated at least 1 hour at room temperature in 10% blocking solution (Roche), followed by overnight incubation at 4°C in 1% blocking solution (10% blocking solution in PBS-t) in which primary antibodies were diluted. Larvae were then washed five times using PBS-t solution and incubated overnight at 4°C in 1% blocking solution containing secondary antibodies. Alexa Fluor 488 goat anti-mouse IgG and Alexa Fluor 564 goat anti-rabbit secondary antibodies (1:2000, Life Technologies) and DAPI diluted 1:1000 in PBS-t were used. Samples were then washed several times in PBS-t solution. Samples were then mounted laterally using 1.5% low-melting point agarose prior to imaging.

Antibody labelling was performed using anti-synaptotagmin2 (znp1) (Developmental Studies Hybridoma Bank, diluted 1:100), anti-FIGQY (neurofascin) (gift from Matthew Rasband, diluted 1:250), anti-detyrosinated tubulin (Life Sciences #RM444, diluted 1:500), anti-acetylated tubulin (Sigma clone 6–11-B-1, diluted 1:1000) and anti-SV2 (Developmental Studies Hybridoma Bank, diluted 1:100).

### Imaging of whole mount fish samples

Injected transgenic fish motoneurons were acquired live at 2 dpf using a 63.3X (W PL APO VIS-IR 421480– 9900), with Z-volumes captured at a stepsize of 0.5 µm, using an Olympus FV3000 laser scanning confocal microscope. Image acquisition was conducted at a resolution of 1024×1024 pixels, with a scan rate of 4 µs/pixel. The same imaging approach was used for the immunostaining experiments on WT and mutant samples fixed at 3 dpf. Image processing and analysis were conducted using ImageJ software.

### Behavioral tests

Free swimming behavioral acquisitions, fish tracking, tail segmentation and analysis were performed as previously described^68^. In total 16 larvae were tested for each genotype with the acquisition lasting for 30 minutes. To assess the seizure susceptibility of our fish, individual larvae were placed at 5 dpf in a 96-well plate. Larvae were left to acclimate for 1 hour prior to recording. Baseline locomotor activity was recorded for 15 min before exposure to pentylenetetrazole (PTZ, Sigma Millipore). Larvae were recorded for 30 min after PTZ addition, at final concentrations of 20 mM or 3 mM. The same camera and illumination setup as in the free-swimming experiment was used, recording at a framerate of 60 Hz. The fish were tracked using the FastTrack software^69,70^. Subsequently, we utilized custom-made Python scripts to extract detailed parameters from the tracking data, including trajectories, speed, distances, latency and to characterize behavioral state.

### Statistical Methods

All comparisons between WT controls and mutant larvae were carried out on Prism 10 (Graphpad). To assess statistical significance the Mann-Whitney test by ranks was performed when the dataset did not follow a normal distribution or did not meet basic t-test assumptions.

## Data availability

Materials can be obtained from E.N., F.D.B., Z.L. and C.J. under a material transfer agreement with the relevant institutions. The structural coordinates for MTCL1 in complex with Taxol-stabilized MTs and under copolymerization conditions have been deposited in the Protein Data Bank (PDB) under accession codes 10GT and 10GY, respectively. Additionally, the cryo-EM maps from this study are available in the Electron Microscopy Data Bank (EMDB) under accession codes EMD-75159 (Taxol MTs - MTCL1^C-MTBD^), EMD-75165 (Copolymerized MTs - MTCL1^C-MTBD^) and EMD-75166 (Copolymerized MTs - MTCL1^C-MTBD^ with Taxol).

**Supplementary Figure 1:**
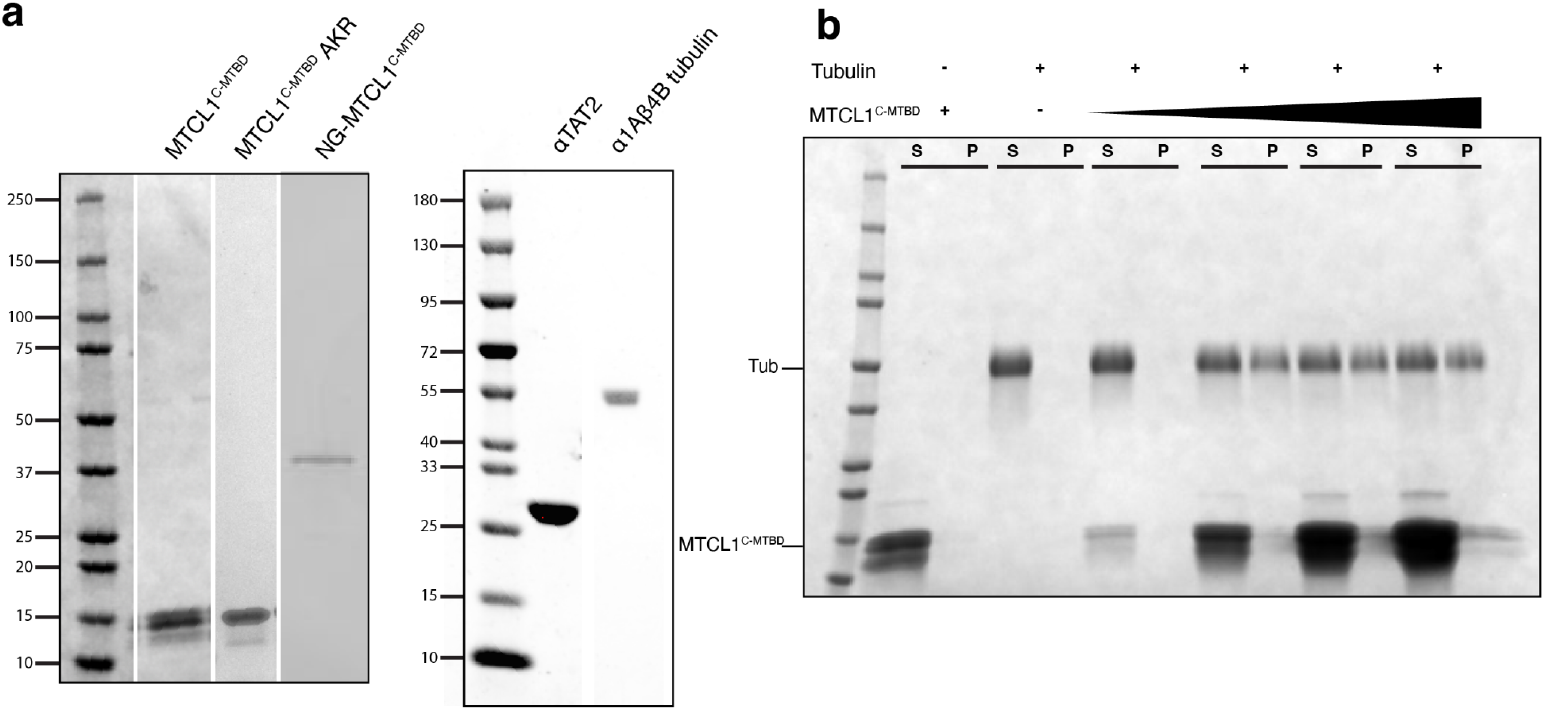
Protein purification and co-sedimentation assay under dynamic MT growth. **(a)** Coomassie Blue staining of the purified proteins used in this study. **(b)** Co-sedimentation assay with tubulin below the critical concentration required for spontaneous *in vitro* MT nucleation. In the absence of MTCL1^C-MTBD^, all tubulin remains in the soluble fraction. In contrast, the addition of MTCL1^C-MTBD^ promotes MT formation, which co-sediments with the MAP.

**Supplementary Figure 2:**
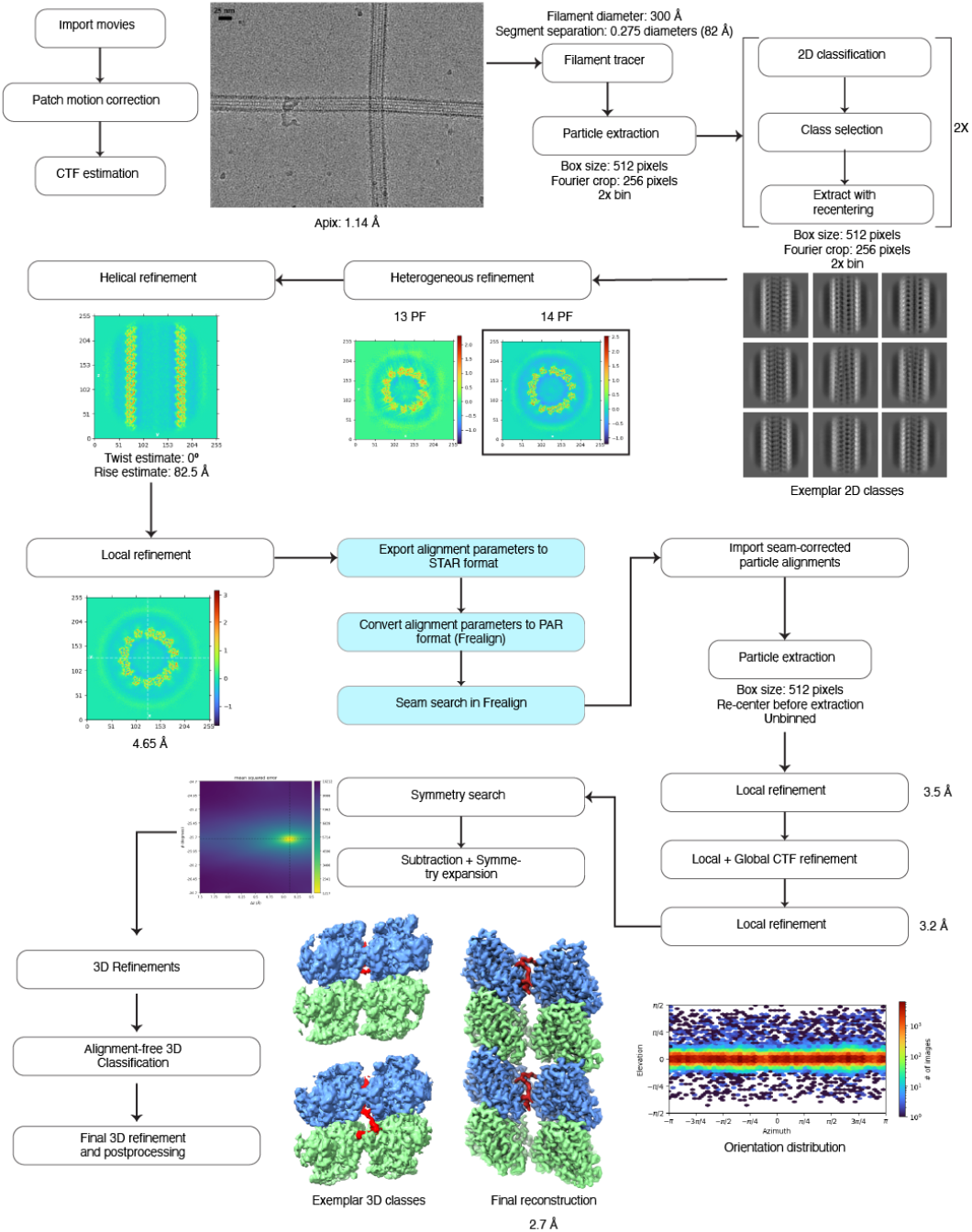
Cryo-EM data processing pipeline for MT-MTCL1^C-MTBD^. Steps highlighted in light blue indicate processes implemented during the seam search protocol.

**Supplementary Figure 3:**
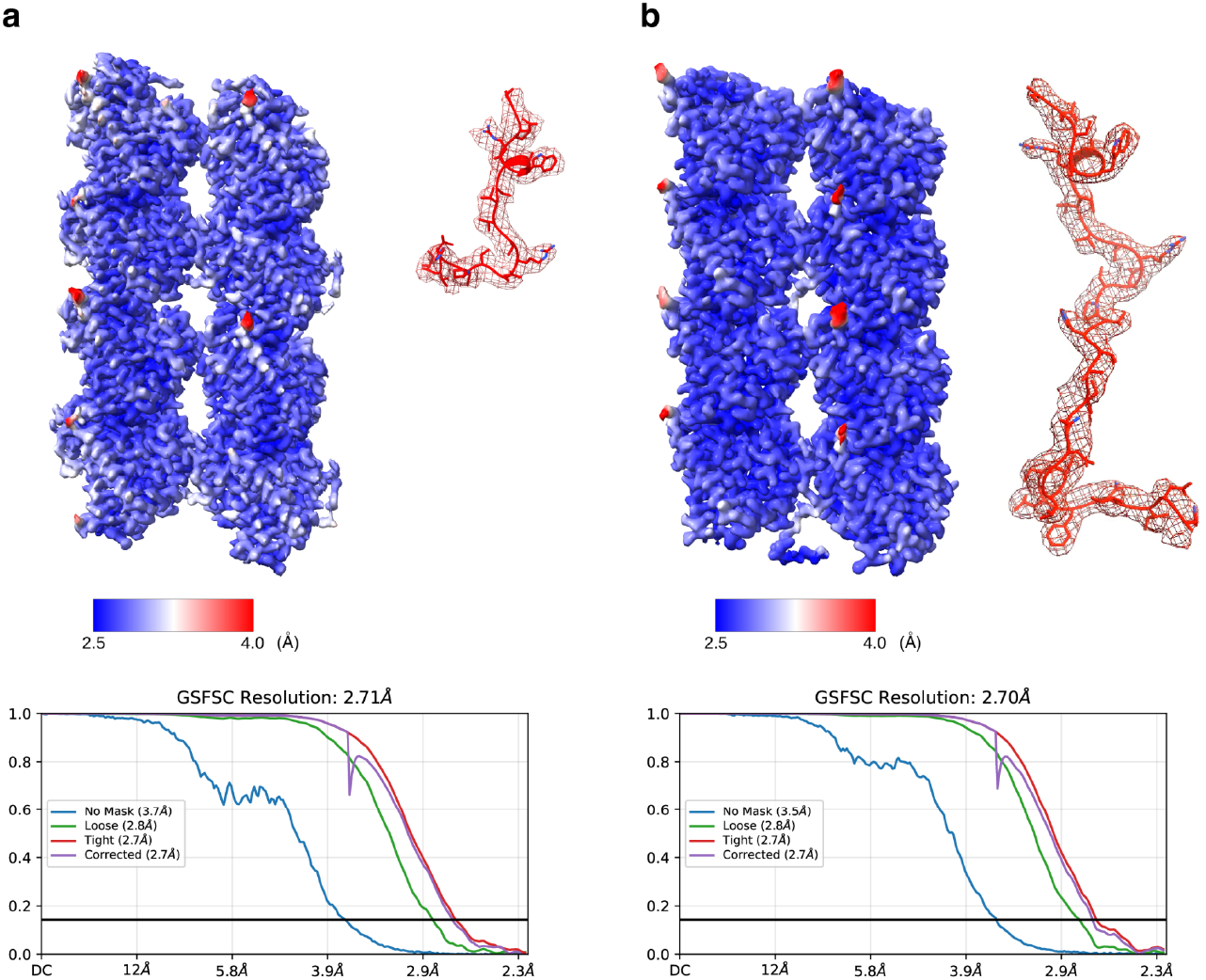
Resolution estimation for the cryo-EM maps of MT-MTCL1^C-MTBD^ complexes. **(a)** Cryo-EM density map colored by local resolution for the MT bound to MTCL1^C-MTBD^ added after assembly (left). Cryo-EM density for MTCL1 shown as a mesh with the refined atomic model fitted inside (right). Gold-standard Fourier shell correlation (FSC) curves indicating global resolution (bottom). **(b)** Cryo-EM density map colored by local resolution for the MT copolymerized with MTCL1^C-MTBD^ (left). Cryo-EM density of MTCL1^C-MTBD^ shown as a mesh with the refined atomic model fitted inside (right). Gold-standard FSC curves indicating global resolution (bottom).

**Supplementary Figure 4:**
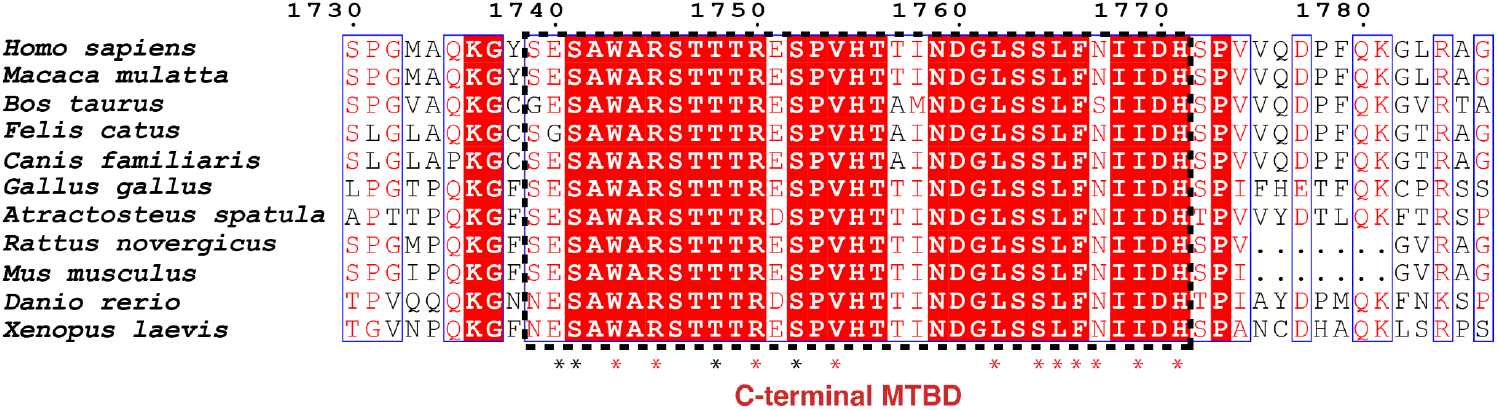
MTCL1 residues that interact with tubulin are conserved across species. Multiple sequence alignment of MTCL1^C-MTBD^ from diverse species, with the human sequence used as the reference. Residues identified as contacting tubulin in our cryo-EM structure through side-chain dependent and side-chain independent contacts are labeled with red and black asterisks, respectively. The figure was generated with ESPript3^71^

**Supplementary Figure 5:**
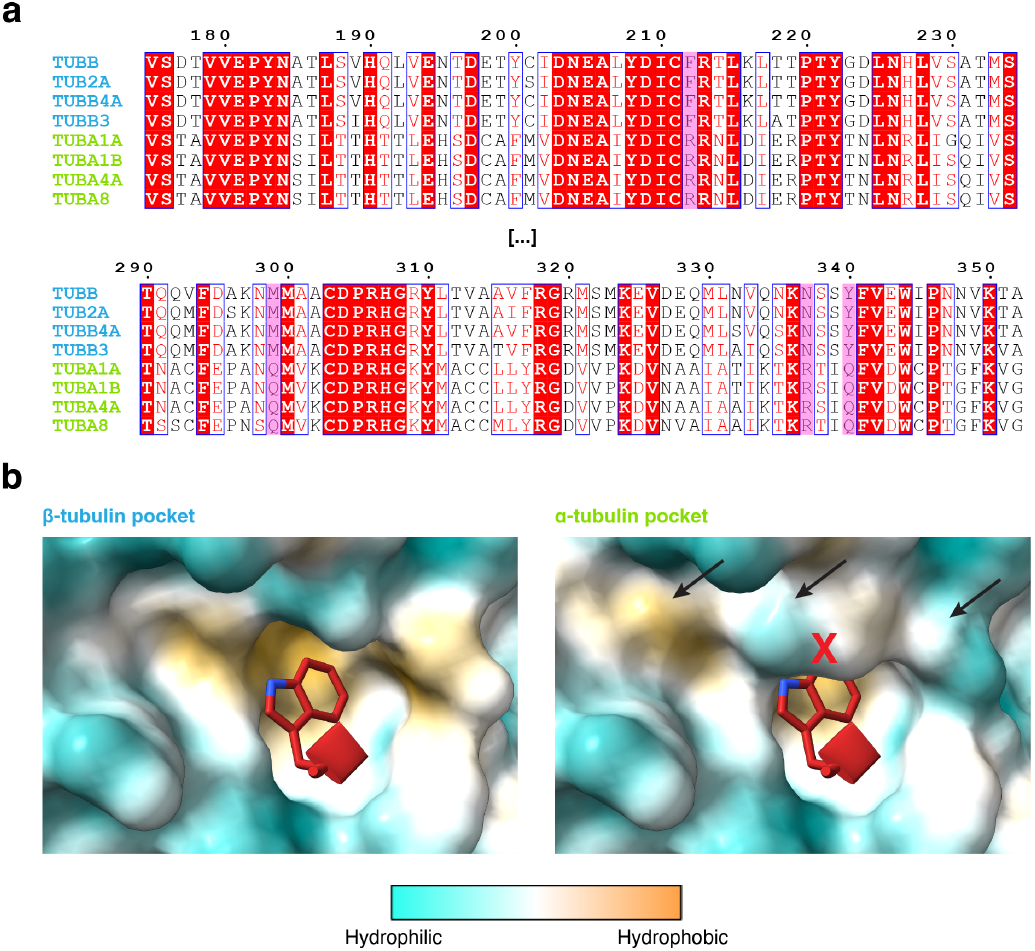
Sequence and structural basis for β-tubulin selectivity of MTCL1^C-MTBD^. **(a)** Sequence alignment of representative pig (Sus scrofa) α- and β-tubulin isotypes highlighting residues that interact with MTCL1^C-MTBD^. The numbering corresponds to the TUBB sequence, which is used as reference. Amino acids that diverge between α- and β-tubulins and contribute to MTCL1 binding are boxed in purple, while fully conserved positions are boxed in red. **(b)** Structural comparison of the W1743-binding pocket in β-tubulin (left) and α-tubulin (right) colored by hydrophobicity. To display the W1743 in the context of the α-tubulin pocket, we used the β-tubulin pocket as a starting point and manually changed the residues in the pocket to those of α-tubulin. Residue substitutions (black arrows) in α-tubulin disrupt the pocket and generate steric clashes (red X).

**Supplementary Figure 6:**
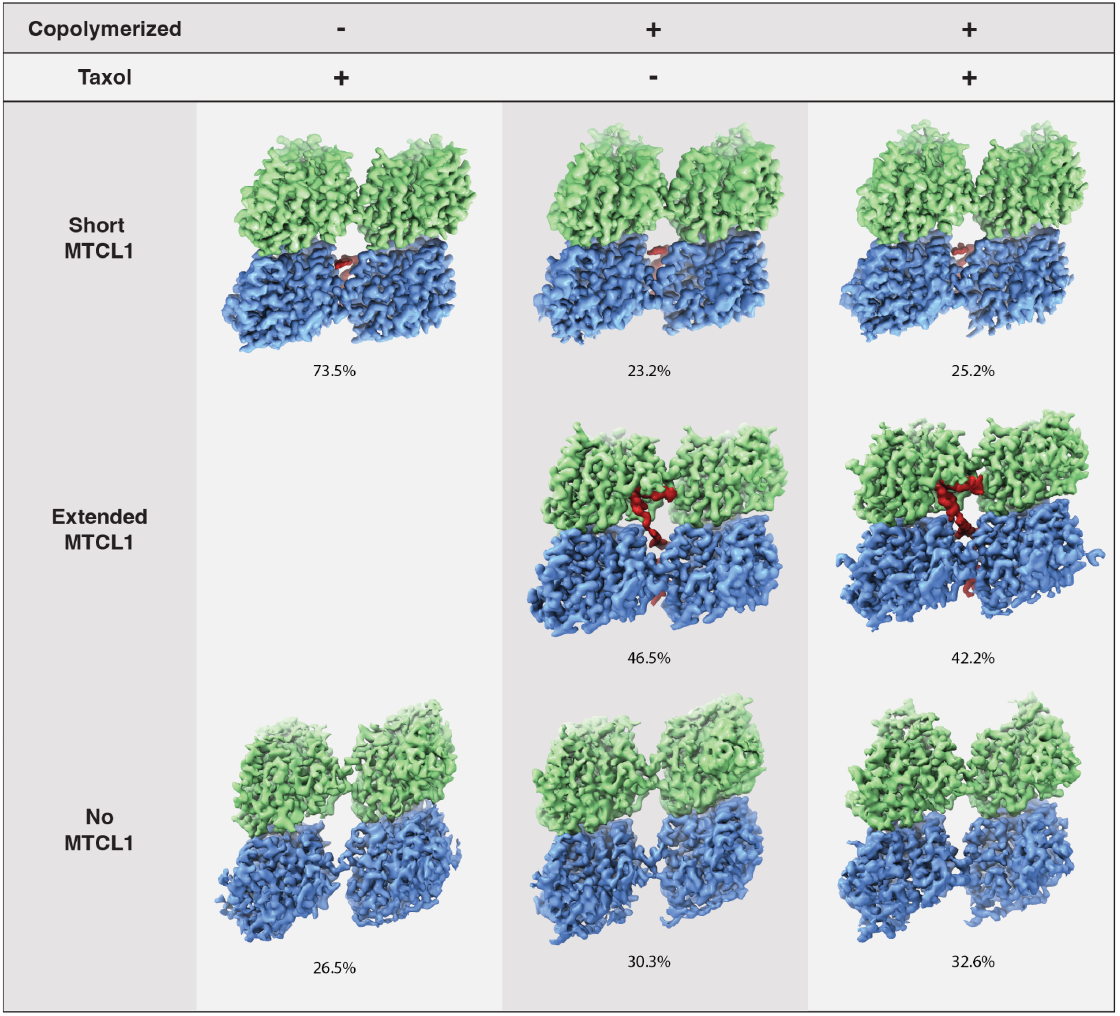
Systematic analysis of cryo-EM populations for three MTCL1C^-MTBD^ datasets. The three datasets correspond to Taxol-stabilized MTs with MTCL1^C-MTBD^ added after polymerization, MTs copolymerized with MTCL1^C-MTBD^ in the absence of Taxol, and MTs copolymerized with MTCL1^C-MTBD^ in the presence of Taxol. For each dataset, reconstructions are shown for tubulin with short MTCL1, extended MTCL1, and no MTCL1, when significantly populated. All classes show a luminal view of a tubulin inter-dimer interface. Percentages denote particle fractions.

**Supplementary Figure 7:**
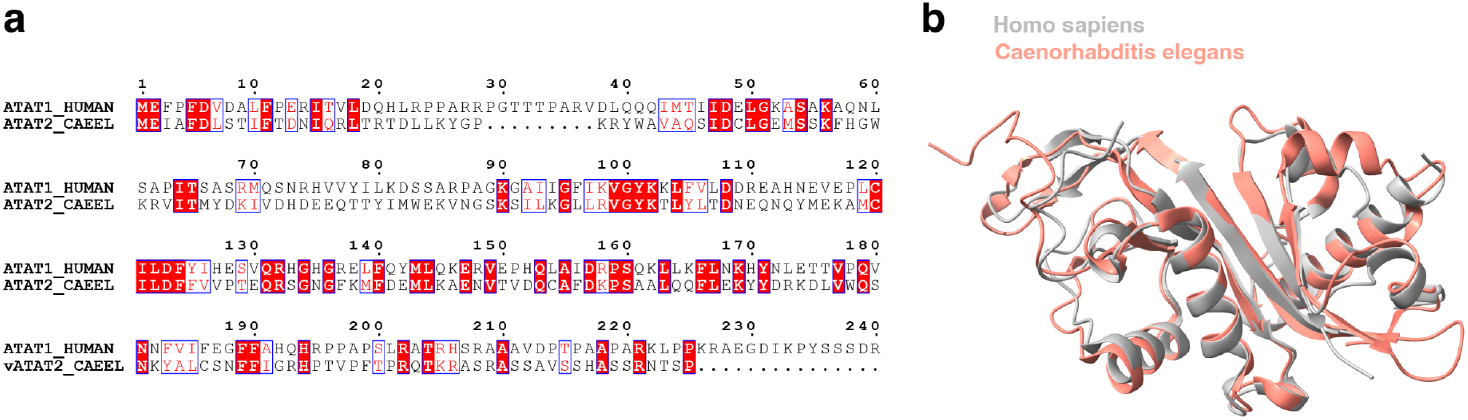
Conservation of human and worm tubulin acetyltransferases. **(a)** Multiple sequence alignment of the catalytic domains of human αTAT1 and *C. elegans* αTAT2. Residues are colored by conservation, with red highlights and red text indicating identity and strong similarity, respectively; numbering corresponds to human αTAT1. **(b)** Structural superposition of the catalytic cores highlighting the conserved GNAT fold. Human αTAT1 (PDB 4GS4) is shown in gray and *C. elegans* αTAT2 (PDB 8Y9F) in salmon. Substantial agreement is observed across the core secondary-structure elements, with variability confined mainly to surface loops.

**Supplementary Figure 8:**
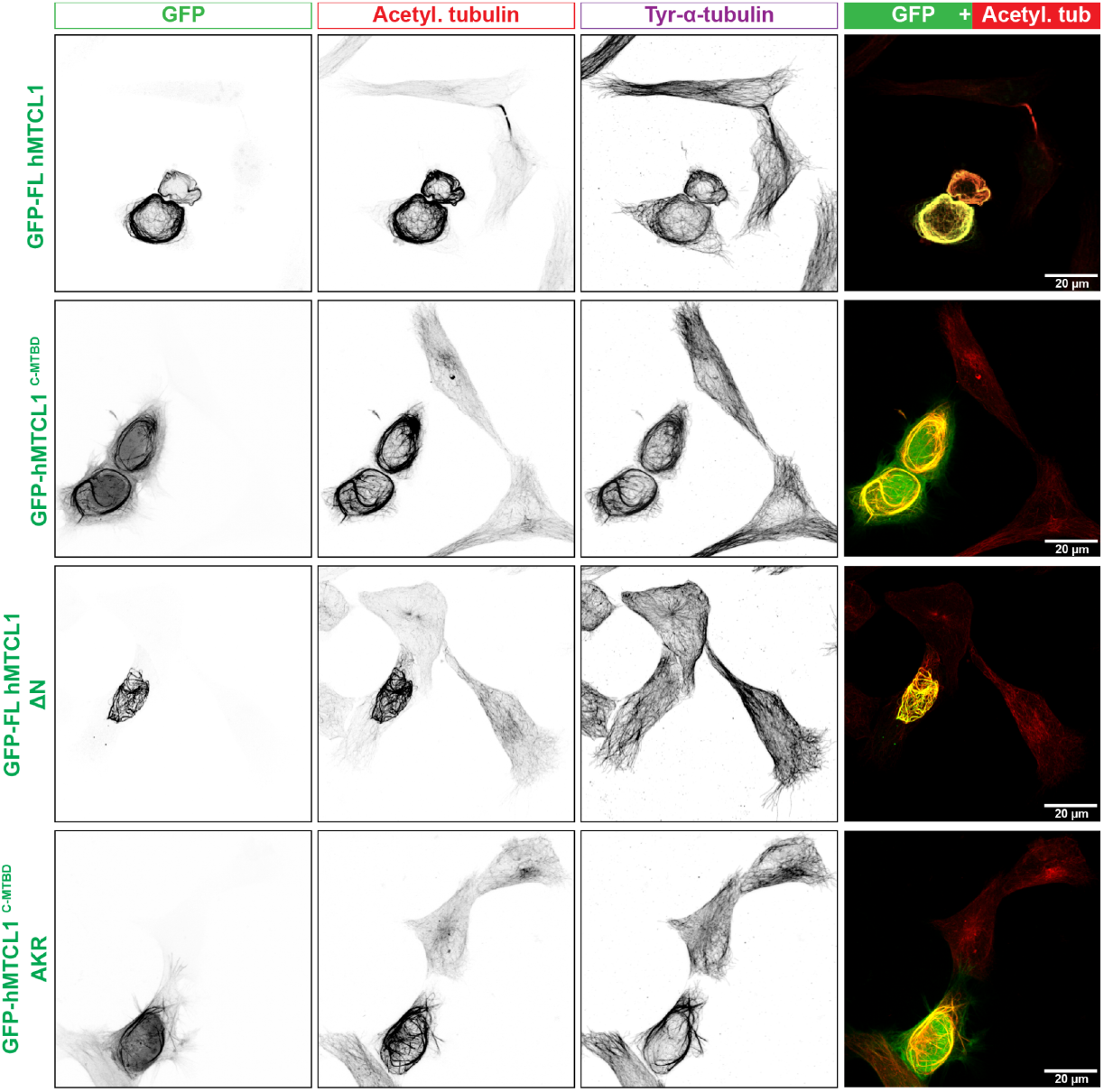
Characterization of α-tubulin K40 acetylation in cells with high levels of MTCL1^C-MTBD^ expression and strong MT bundling. Representative immunofluorescence images of HeLa cells expressing GFP-tagged human MTCL1 plasmids for 24 hours. The images focus on cells with strong expression levels and pronounced MT bundling. Cells were stained with anti-α-tubulin acetylation and tyrosination antibodies. Scale bar, 20 μm. Number of cells analyzed from three independent experiments: GFP-FL hMTCL1, n = 13 cells; GFP-hMTCL1^C-MTBD^, n = 13 cells; GFP-FL hMTCL1 ΔN, n = 13 cells; GFP-hMTCL1^C-MTBD^ AKR, n = 15 cells.

**Supplementary Figure 9:**
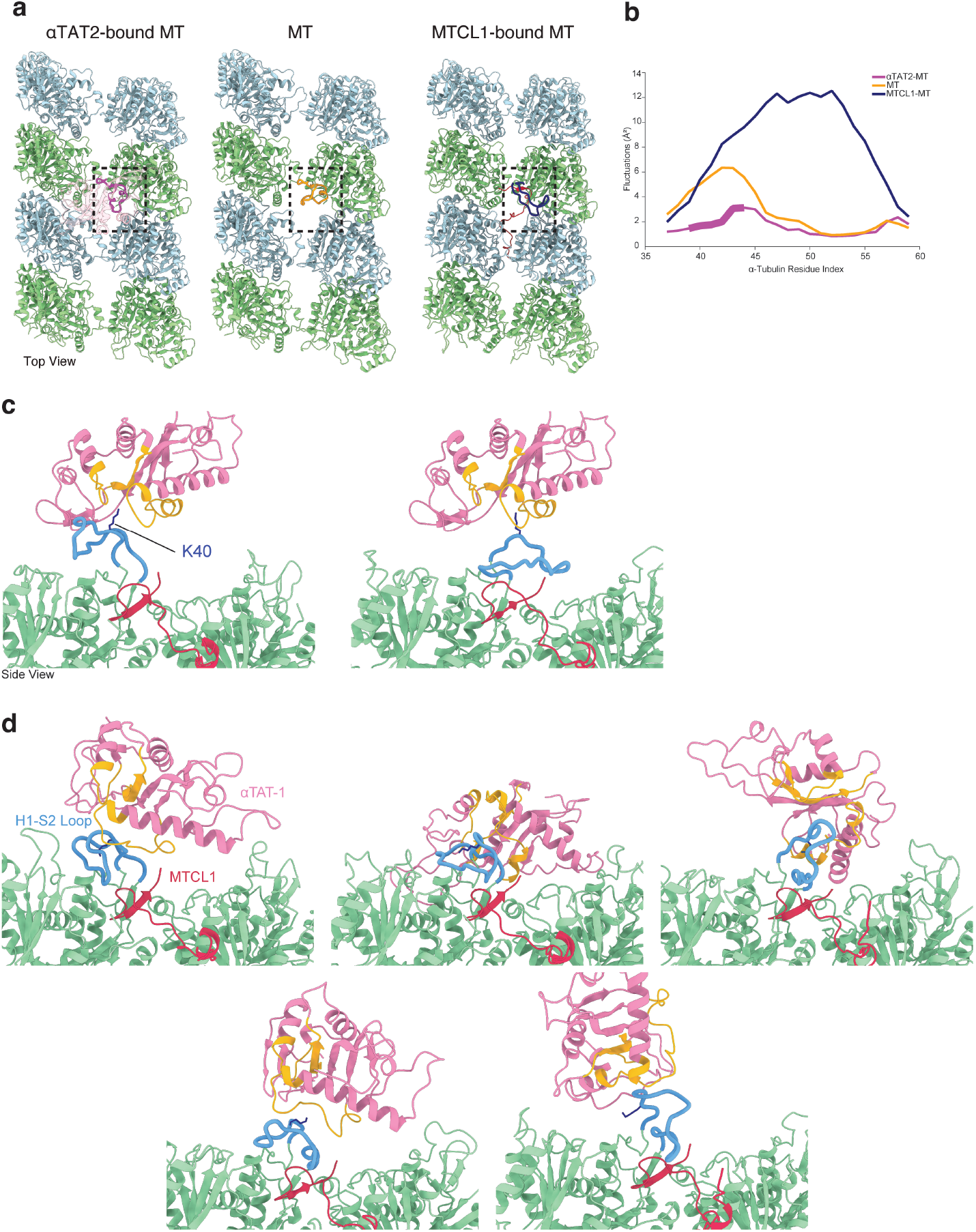
MTCL1^C-MTBD^ increases H1–S2 loop flexibility and promotes αTAT1 capture in all-atom MD simulations. **(a)** Representative all-atom MD simulations snapshots of the H1–S2 loop in the αTAT2-bound MT (left), in the apo MT after αTAT2 removal (middle), and in the presence of MTCL1^C-MTBD^ (right). All MTs are shown in a luminal view. **(b)** Per-residue Cα residue fluctuation profiles for the H1–S2 loop in the three simulation conditions described in a. Residues that adopt a structured conformation in the αTAT2–bound state are highlighted in bold in the residue fluctuation plot. **(c)** Starting conformations used to initiate the two MD simulations of the MTCL1^C-MTBD^–tubulin–αTAT1 ternary complex shown in Figure 3h. In each run, αTAT1–acetyl-CoA was placed ∼10 Å from the tubulin surface and the H1–S2 loop was initialized in a distinct conformation. **(d)** Additional αTAT1 binding poses observed in ternary-complex MD simulations. Representative bound conformations of αTAT1 on the MT surface in the MTCL1^C-MTBD^–tubulin–αTAT1 ternary complex from five independent simulations initiated with different starting H1–S2 loop conformations (for a total of seven independent simulations, including those shown in Figure 3h), showing stable association of αTAT1 and interaction with the H1–S2 loop. In c and d, residues corresponding to the αTAT1 active site are shown in orange.

**Supplementary Figure 10:**
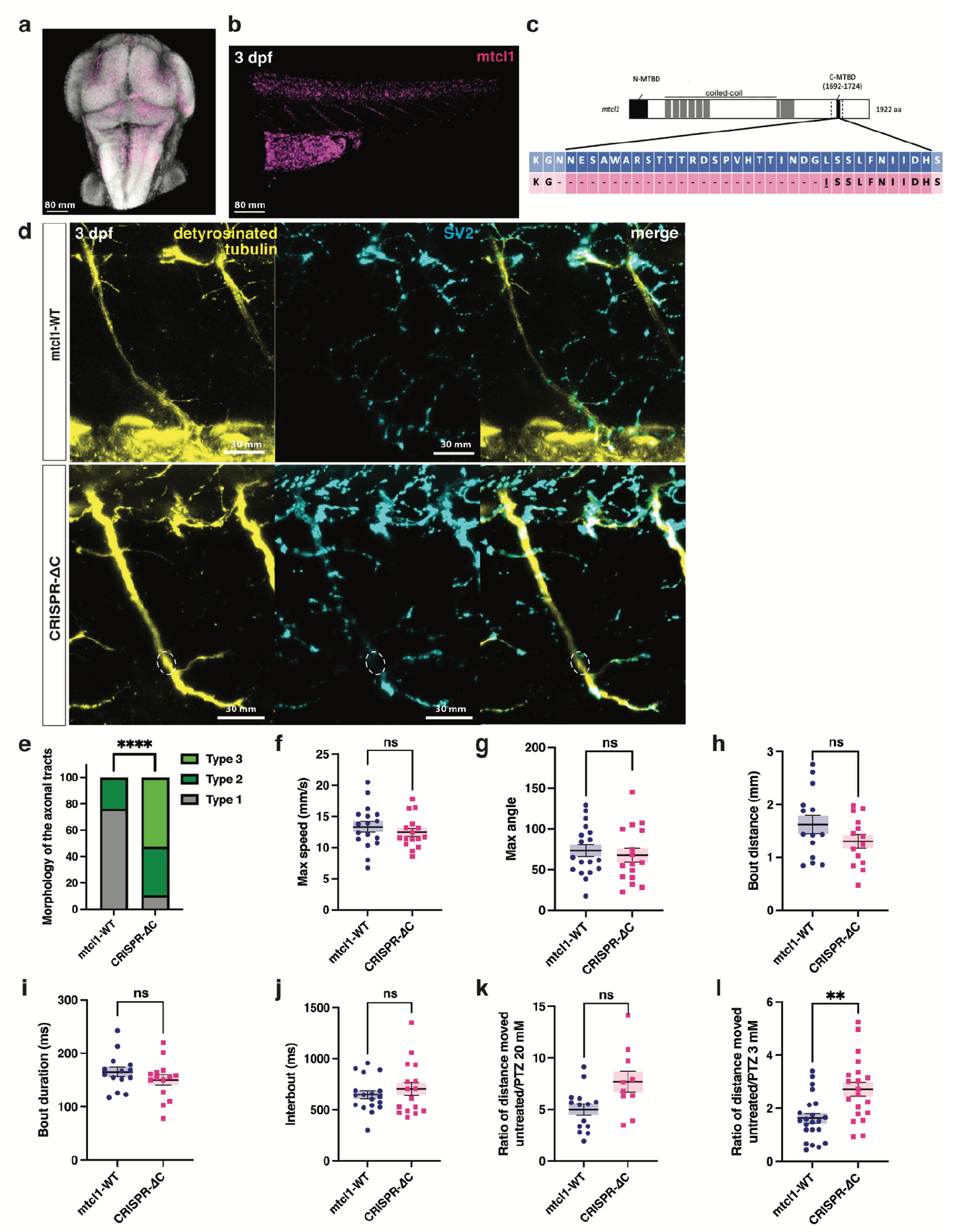
Characterization of *mtcl1* in zebrafish. **(a, b)** Confocal Z-stack images of a 3 dpf WT zebrafish brain (a) and spine (b) stained by multiplex HCR *in situ* hybridization show *mtcl1* transcripts (magenta; hairpin fluorophore 594 nm) expressed in the whole zebrafish CNS. DAPI (gray) marks cell nuclei in the brain. **(c)** Domain architecture of zebrafish *mtcl1* with CRISPR-Cas9-induced in-frame deletion in pink. **(d)** Developing CaP neurons labelled for detyrosinated tubulin (yellow) from *mtcl1-WT* and *CRISPR-ΔC* mutant zebrafish were co-labelled for SV2 (cyan) at 3 dpf. Swellings are highlighted with dotted white circles. Scale bar = 30 mm. **(e)** Percentage of types 1, 2 and 3 axonal morphologies in 3 dpf control and mutant zebrafish labelled for acetylated tubulin (*mtcl1-WT*, n = 21; *CRISPR-ΔC*, n = 19; \*\*\*\**p < 0*.*0001*). Data represent mean and SEM. Statistical significance was assessed by a Mann-Whitney U-test. **(f-j)** Behavioral characterization of *mtcl1 CRISPR-ΔC* mutants. Tracking of 6 dpf larvae freely swimming in a Petri dish does not show any difference between *mtcl1-WT* and *CRISPR-ΔC* mutant larvae for max speed (f), max tail angle (g), bout distance (h), bout duration (i) and interbout (j) (*mtcl1-WT*, n = 17; *CRISPR-ΔC*, n = 16). Data represent mean and SEM. **(k, l)** At 6 dpf, larval zebrafish locomotor behavior recorded for 15 min before and after PTZ exposure, allowed us to compare a ratio between distance moved before and after PTZ exposure. 3 mM PTZ (k) (*mtcl1-WT*, n = 22; *CRISPR-ΔC*, n = 20) or 20 mM PTZ (l) (*mtcl1-WT*, n = 14; *CRISPR-ΔC*, n = 11). Data represent mean and SEM. Statistical significance was determined by ANOVA (*p* = 0.121 for PTZ 20mM, \*\**p* = 0.002 for PTZ 3mM).

**Supplementary Table 1:**
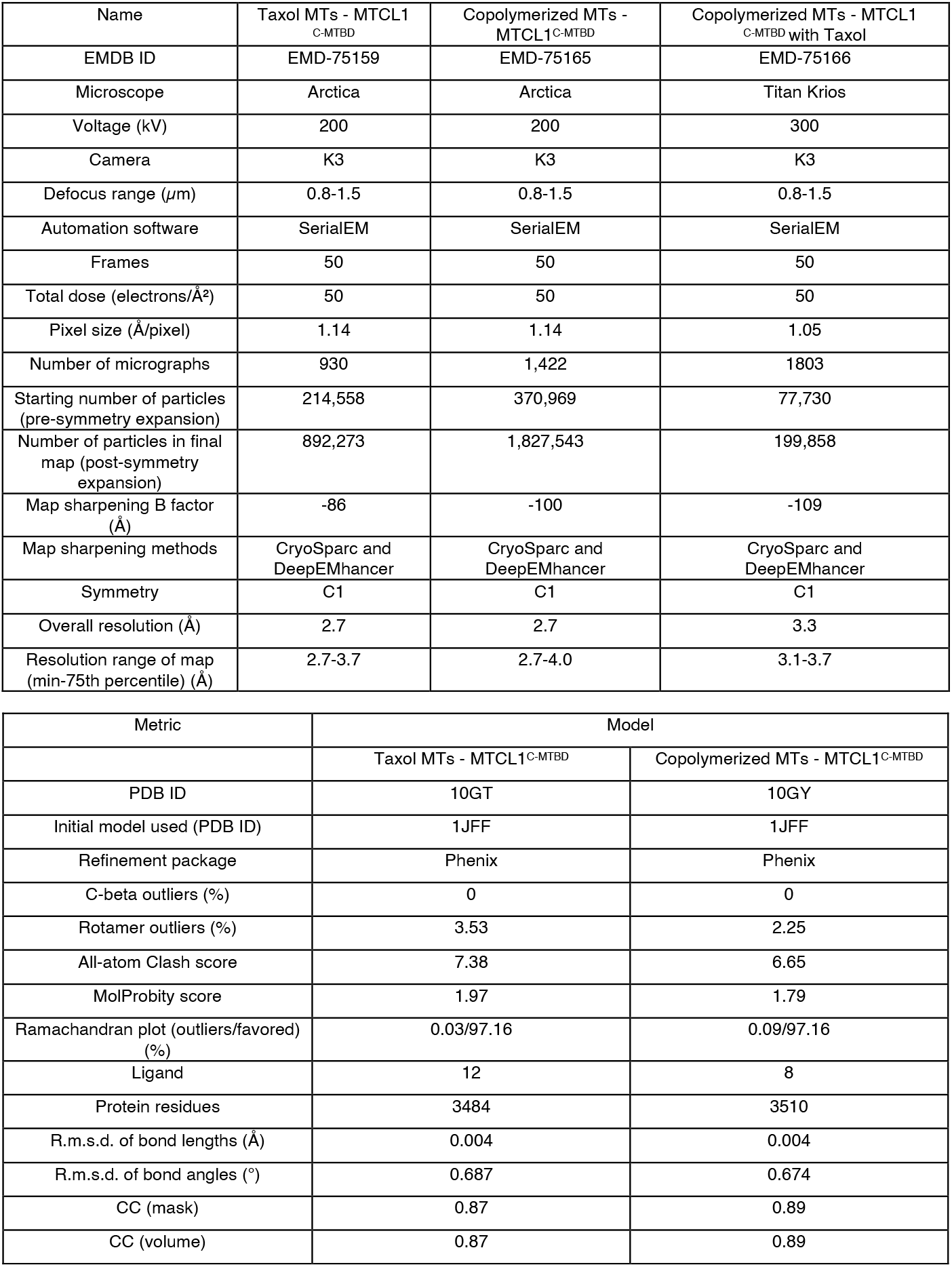
Cryo-EM data collection and model real-space refinement parameters.

**Supplementary Table 2:** The list of amino acids from α-tubulin and β-tubulin whose C_α_ atoms were constrained during MD simulations.

| Tubulin | Constrained Residues |
| --- | --- |
| α-Tubulin | T382-S439 |
| β-Tubulin | T372-D427 |

